# p38-mitogen activated kinases mediate a developmental regulatory response to amino acid depletion and associated oxidative stress in mouse blastocyst embryos

**DOI:** 10.1101/807305

**Authors:** Pablo Bora, Vasanth Thamodaran, Andrej Šušor, Alexander W. Bruce

## Abstract

Maternal starvation coincident with preimplantation development has profound consequences for placental-foetal development, with various identified pathologies persisting/manifest in adulthood; the ‘Developmental Origin of Health and Disease’ (DOHaD) hypothesis/model. Despite evidence describing DOHaD-related incidence, supporting mechanistic and molecular data relating to preimplantation embryos themselves are comparatively meagre. We recently identified the classically recognised stress-related p38-mitogen activated kinases (p38-MAPK) as regulating formation of the extraembryonic primitive endoderm (PrE) lineage within mouse blastocyst inner cell mass (ICM). Thus, we wanted to assay if PrE differentiation is sensitive to amino acid availability, in a manner regulated by p38-MAPK. Although blastocysts appropriately mature, without developmental/morphological or cell fate defects, irrespective of amino acid supplementation status, we found the extent of p38-MAPK inhibition induced phenotypes was more severe in the absence of amino acid supplementation. Specifically, both PrE and epiblast (EPI) ICM progenitor populations remained unspecified and there were fewer cells and smaller blastocyst cavities. Such phenotypes could be ameliorated, to resemble those observed in groups supplemented with amino acids, by addition of the anti-oxidant NAC (N-acetyl-cysteine), although PrE differentiation deficits remained. Therefore, p38-MAPK performs a hitherto unrecognised homeostatic early developmental regulatory role (in addition to direct specification of PrE), by buffering blastocyst cell number and ICM cell lineage specification (relating to EPI) in response to amino acid availability, partly by counteracting induced oxidative stress; with clear implications for the DOHaD model.

## Introduction

The formation of the peri-implantation stage mouse blastocyst (by embryonic day 4.5/E4.5) represents the culmination of the preimplantation period in which three distinct cell lineages emerge. Two lineages are differentiating and will ultimately yield supportive extraembryonic tissues; the outer-residing and epithelised trophectoderm (TE) that gives rise to the placenta, and the mono-layered primitive endoderm (PrE), occupying the interface between the fluid filled cavity and underlying inner cell mass (ICM), that contributes to the yolk sac. The third lineage, represented by the pluripotent epiblast (EPI), residing deep within the ICM, serves as a progenitor pool for all subsequent tissues of the foetus; extensively reviewed (Frum and Ralston, 2015;Chazaud and Yamanaka, 2016;Rossant, 2016)). We have previously reported a role for p38-mitogen activated kinases α/β(herein referred to as p38-MAPK), employing pharmacological inhibition, in regulating primitive endoderm (PrE) differentiation within mouse blastocyst inner cell mass (ICM)(Thamodaran and Bruce, 2016). p38-MAPK was found to act during the early stages of ICM maturation (the period between E3.5-E4.5), downstream of fibroblast growth factor (FGF) signalling, permitting PrE progenitors to resolve their uncommitted fate (as is characteristic of the majority of nascent ICM cells at E3.5 (Chazaud et al., 2006)) and thus, functionally diverge and segregate from the EPI lineage (Thamodaran and Bruce, 2016). Although, p38-MAPK (and their related paralogs, p38γ/δ) belong to the wider family of serine-threonine and tyrosine kinases, regulating a wide variety of cellular functions (Cargnello and Roux, 2011), they differ from the other family members, such as extra-cellular regulated kinases [*e.g.* ERK1/2 – themselves implicated in FGF-mediated PrE differentiation at a developmental point succeeding that identified for p38-MAPK (Nichols et al., 2009;Yamanaka et al., 2010;Frankenberg et al., 2011;Kang et al., 2013;Thamodaran and Bruce, 2016)] in that they are classically known to be activated by extracellular stress stimuli; for example pro-inflammatory cytokines, U.V. radiation and physical stress, rather than liganded growth-factor associated receptor tyrosine kinases (Remy et al., 2010). It is estimated activated p38-mitogen activated kinases in general are able to phosphorylate and regulate between 200-300 cellular substrates (Cuadrado and Nebreda, 2010;Trempolec et al., 2013;Hornbeck et al., 2019). In the context of this study, there is precedent for the involvement of active p38-MAPK in amino acid (AA) signalling (Casas-Terradellas et al., 2008) and regulation of autophagy (Webber, 2010), in other non-embryo models.

The ‘Developmental Origin of Health and Disease’ (DOHaD) model hypothesises environmental cues, particularly nutrient availability, during the peri-conceptual days of development manifests changes in embryonic metabolism and development, with potential pathological consequences extending into adulthood (O’Brien, 1999). Rodent pups born to mothers exposed to low protein diets during preimplantation stages of development are reported to have significantly increased birth weight, elevated systolic blood pressure, liver hypertrophy, cardiovascular and metabolic disorders, aberrant establishment of gene imprints plus hyperactive behaviour and poor memory (Kwong et al., 2000;Kwong et al., 2006;Fleming et al., 2015;Fleming et al., 2018). Such observations are indicative of adaptive and persistent changes in early embryo physiology/metabolism; indeed, similar studies report increased endocytosis and nutrient uptake in the extraembryonic TE and PrE lineages, arising from altered epigenetic gene regulation (Sun et al., 2014;Sun et al., 2015). Thus, mounting evidence corroborates the incidence of the DOHaD model, yet there is a comparative dearth of detailed and supportive molecular mechanistic data that could underpin how preimplantation stage embryos react and develop under conditions of nutrient deprivation.

Accordingly, we have examined our previously identified p38-MAPK inhibition induced defective PrE phenotypes, themselves associated with reduced blastocyst cell number (Thamodaran and Bruce, 2016), to ascertain if p38-MAPK not only regulates PrE differentiation *per se*, but also perform a dual regulative/homeostatic role in mediating preimplantation mouse embryo/blastocyst development in response to limited amino acid (AA) availability, as could be inferred from earlier studies (Kwong et al., 2000;Kwong et al., 2006;Sun et al., 2014;Fleming et al., 2015;Sun et al., 2015;Fleming et al., 2018) addressing DOHaD.

## Materials & Methods

### Mouse lines and embryo culture

All experimental procedures relating to mice (*i.e.* derivation of preimplantation stage embryos for further study) complied with ‘ARRIVE’ guidelines and were carried out in accordance with EU Directive 2010/63/EU (for animal experiments). Superovulation and strain mating regime to produce embryos for the experiments are shown in the figures and are as previously described (Mihajlovic et al., 2015). E1.5 (*i.e.* 2-cell) stage embryos were isolated from the oviducts of the females in M2 media (pre-warmed at 37°C for at least 2-3 hours) and thereafter cultured in KSOM (EmbryoMax® KSOM Mouse Embryo Media; cat. # MR-020P-5F - pre-warmed and equilibrated in 5% CO_2_ and 37°C), either with or without amino acid (AA) supplementation. For KSOM+AA condition, Gibco™ MEM Non-Essential Amino Acids Solution (100×) (cat. # 11140035) and Gibco™ MEM Amino Acids Solution (50×) (cat. # 11130036) were used to a working concentration of 0.5×. Embryos were cultured in micro-drops prepared in 35mm tissue culture dishes covered with light mineral oil (Irvine Scientific. cat. # 9305), in 5% CO_2_ incubators maintained at 37°C until the appropriate stage and thereafter were analysed according to the experimental design. Chemical inhibition of p38-MAPKs was carried out using SB220025 (Calbiochem® cat. # 559396; dissolved in dimethyl sulfoxide/DMSO) at 20μM working concentration in the respective culture medium, as described previously (Thamodaran and Bruce, 2016). DMSO (Sigma-Aldrich® cat. # D4540) of equivalent volume was used as solvent control to a final working concentration of 0.2% by volume. Embryos with a blastocoel cavity occupying approximately 50% of the volume of the embryo at 12.00 hours on E3.5 were moved to either inhibitory or control culture conditions and cultured for a further 24 hours (*i.e.* E4.5). Rescue experiments were similarly performed by further addition of N-Acetyl-L-cysteine (NAC, dissolved in water; Sigma-Aldrich® cat. # A7250) to a final working concentration of 1 or 10mM in respective culture medium. All KSOM based culture media, with or without additional chemicals (AAs, inhibitors or anti-oxidants), was pre-warmed and equilibrated in 5% CO_2_ and 37°C for at least 3-4 hours prior to embryo transfer.

### Bright-field microscopy, immunofluorescence staining, confocal microscopy and image analysis

Bright-field images were captured using Olympus IX71 inverted fluorescence microscope and Optika TCB3.0 imaging unit along with the associated Optika Vision Lite 2.1 software. To remove the *zona pellucida*, blastocysts were quickly washed and pipetted in pre-warmed drops of Tyrode’s Solution, Acidic (Sigma-Aldrich® cat. # T1788), until the *zona* was visually undetectable, immediately followed by washes through pre-warmed drops of M2 media. Thereafter embryos were fixed, in the dark, at stages with 4% paraformaldehyde (Santa Cruz Biotechnology, Inc. cat. # sc-281692) for 20 minutes at room temperature. Permeabilisation was performed by transferring embryos to a 0.5% solution of Triton X-100 (Sigma-Aldrich® cat. # T8787), in phosphate buffered saline (PBS), for 20 minutes at room temperature. Washes post-fixation, permeabilisation and antibody staining were performed in PBS with 0.05% of TWEEN® 20 (Sigma-Aldrich® cat. # P9416) (PBST) by transferring embryos between two drops or wells (of 96-well micro-titre plates) of PBST, for 20 minutes at room temperature. Blocking and antibody staining was performed in 3% bovine serum albumin (BSA; Sigma-Aldrich® cat. # A7906) in PBST. Blocking incubations of 30 minutes at 4°C were performed before both primary and secondary antibody staining; primary antibody staining (in blocking buffer) was incubated overnight (~16 hours) at 4°C and secondary antibody staining carried out in the dark at room temperature for 70 minutes. Stained embryos were mounted in DAPI containing mounting medium VECTASHIELD® (Vector Laboratories, Inc. cat. # H-1200), placed on cover slips and incubated at 4°C for 30 minutes in the dark, prior to confocal imaging. Details of the primary and secondary antibody combinations used can be found in the supplementary information (table S4). Confocal images were acquired using a FV10i Confocal Laser Scanning Microscope and FV10i-SW image acquisition software (Olympus®). Images were analysed using FV10-ASW 4.2 Viewer (Olympus®) and Imaris X64 Microscopy Image Analysis Software (version 6.2.1; Bitplane AG (Oxford Instruments plc). Cells were counted manually and automatically using Imaris X64.

### Cell number quantification, statistics and graphical representation

Total cell number counts (based on DAPI nuclei staining) were further sub categorised as EPI or PrE cells based on detectable and exclusive NANOG and GATA4 (confocal images in figure 1 and graphs in figure 2, 4 and 5) or GATA6 (confocal images and graphs in figure 5) double immuno-staining, respectively. Cells not located within blastocyst ICMs that also did not stain for either GATA4 and/or NANOG, were designated as outer/ TE cells. Specifically relating to figure 5, ICM cells that were positively stained for both GATA6 and NANOG at E4.5 were designated as uncommitted in terms of cell fate. Initial recording and data accumulation was carried out using Microsoft Excel and further statistical analysis and graphical representations performed with GraphPad Prism 8. A Mann-Whitney pairwise statistical test was employed. Unless otherwise stated within individual graphs as a specific P value (if statistically insignificant), the stated significance intervals were depicted as such: P value < 0.0001 (****), 0.0001 to 0.001 (***), 0.001 to 0.01 (**) and 0.01 to 0.05 (*). All graphs represent dot plots of the total sample size, with associated means and the standard error bars highlighted. Supplementary table S5 summarises the results of additional Welch’s ANOVA tests performed across DMSO and p38-MAPK inhibited conditions in which either no NAC, 1mM NAC and 10mM NAC, was supplemented to both KSOM and KSOM+AA conditions. Analysis was performed using GraphPad Prism 8, P values are numerically stated, and significance intervals are depicted as such: P value < 0.0001 (****), 0.0001 to 0.001 (***), 0.001 to 0.01 (**) and 0.01 to 0.05 (*).

**Figure 1 (colour).**
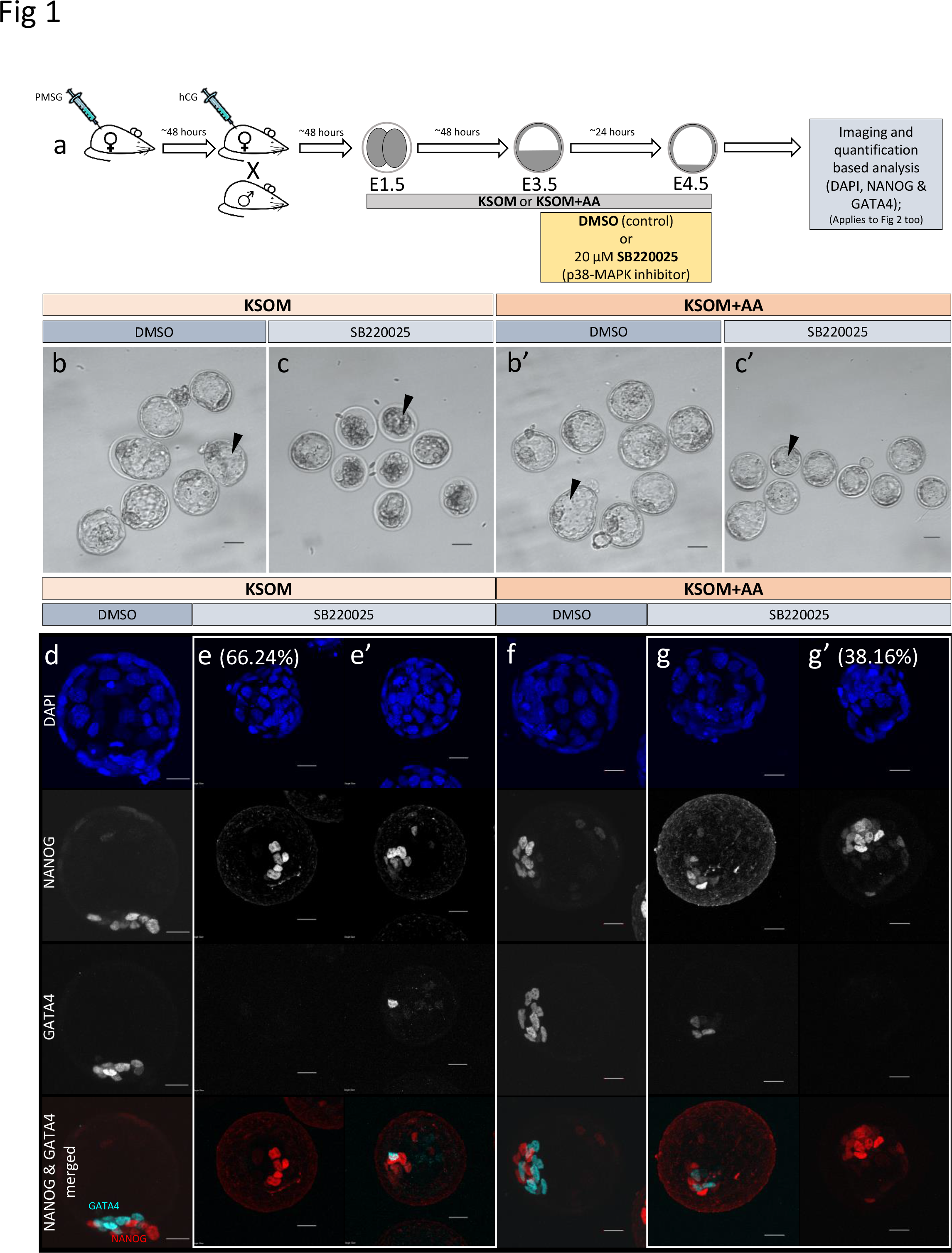
Effect of p38-MAPK inhibition on mouse blastocyst morphology and ICM cell fate derivation in culture conditions ± exogenous amino acid supplementation. **a.** Experimental design: Embryos were collected at E1.5 (2-cell stage) and *in vitro* cultured to E3.5 in media without (KSOM) or with amino acid supplementation (KSOM+AA) and transferred to respective control (DMSO) or p38-MAPK inhibitory conditions (SB220025) until E4.5. Embryos were then fixed, immuno-stained and imaged as described in materials and methods. **b to c’.** Bright-field micrographs of mouse blastocysts at E4.5; all treatments were carried out from E3.5 to E4.5 *i.e.* 24 hours. Panels, from left to right, represent KSOM + DMSO (b), KSOM + p38-MAPK inhibition (c), KSOM+AA + DMSO (b’) and KSOM+AA + p38-MAPK inhibition (c’). Black arrowheads notify presence, absence and relative volumes of the blastocyst cavities. In KSOM + p38-MAPK inhibition (c), blastocoel cavities are markedly smaller and/or collapsed, whereas mostly intact cavities are observed in all other conditions. Scale bar = 40μm. **d to g’.** Z-stack projection confocal images of embryos at E4.5 under the conditions and treatments as depicted above/ panel (a); stained for, from top to bottom, nucleus/ DNA (DAPI), epiblast (NANOG), primitive endoderm (GATA4) and total inner cell mass (NANOG and GATA4 merged) under, KSOM + DMSO (d), KSOM + p38-MAPK inhibition, with no GATA4 positive cells (66% of analysed embryos) (e) and with one or more GATA4 positive cells (34% of analysed embryos) (e’), KSOM+AA + DMSO (f) and KSOM+AA + p38-MAPK inhibition with one or more GATA4 positive cells (65% of analysed embryos) (g) and with no GATA4 positive cells (35% of analysed embryos) (g’) culture conditions. All images in one vertical panel are of the same embryo at the same magnification. Scale bar = 20μm.

**Figure 2 (colour).**
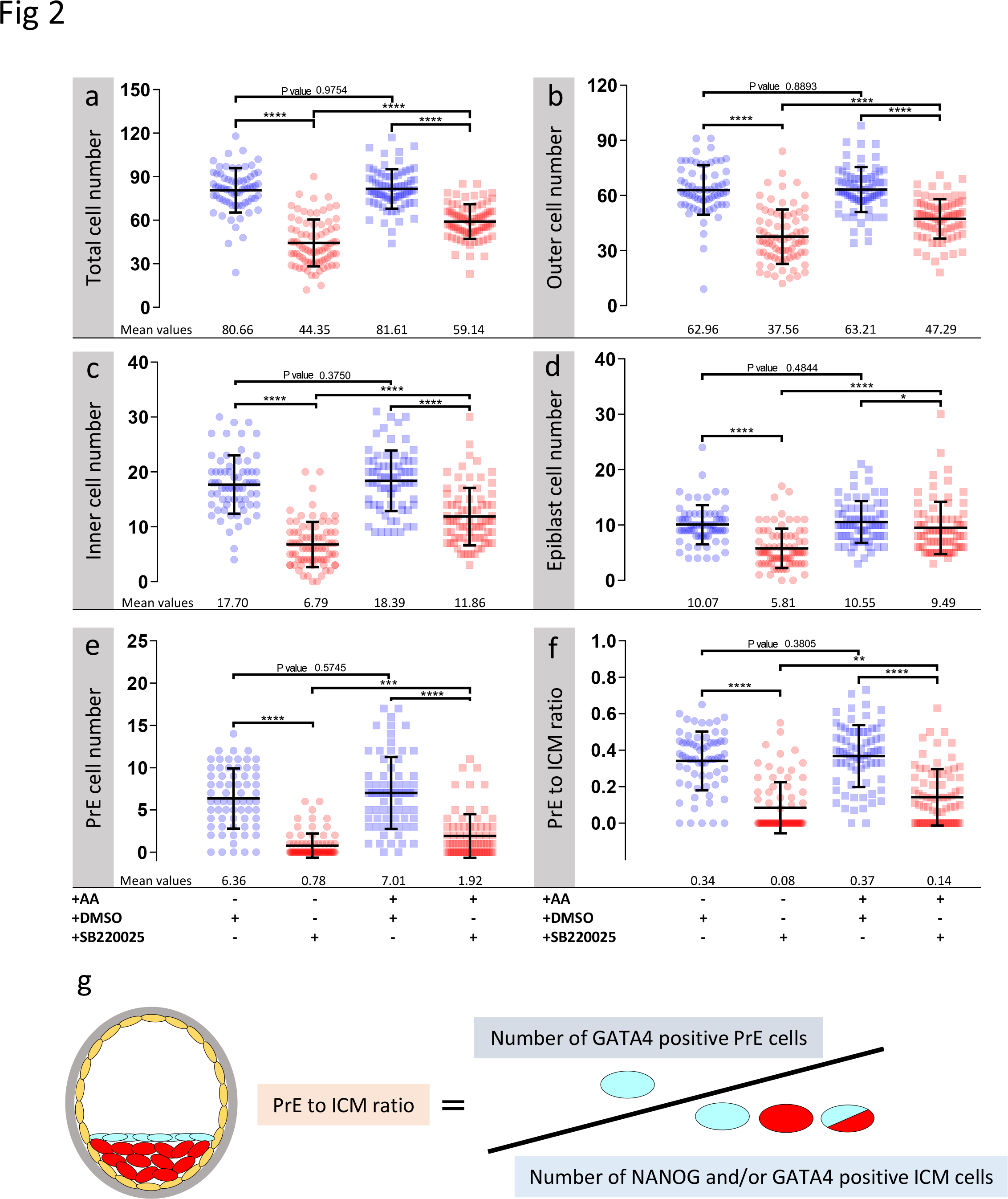
Effect of p38-MAPK inhibition on blastocyst cell numbers, based on DAPI, NANOG and GATA4 staining and confocal microscopy, in culture conditions ± exogenous amino acid supplementation. Culture conditions in relation to amino acid supplementation status and presence of DMSO or p38-MAPK inhibitor (SB220025), from E3.5-E4.5 (as shown Fig. 1) are also shown at the bottom of the individual charts; from left to right; KSOM + DMSO (n = 67), KSOM + p38i (n = 77), KSOM+AA + DMSO (n = 71) and KSOM+AA + p38i (n = 76). **a.** Quantification of total cell number based on counting DAPI stained blastomere nuclei. **b.** Quantification of outer cell number based on subtracting NANOG and/or GATA4 positive inner cells from the DAPI stained total cell number (plus position within the embryo). **c.** Quantification of inner cell number based on NANOG and/or GATA4 stained inner cells (plus position within the embryo). **d.** Quantification of epiblast (EPI) cell number based on NANOG alone stained inner cells. **e.** Quantification of primitive endoderm (PrE) cell number based on GATA4 alone stained inner cells. **f.** Contribution of GATA4 stained primitive endoderm cells as a ratio of the total inner cells (NANOG and/or GATA4 stained cells). **g.** Explanatory scheme of method employed to calculate PrE:ICM ratio values depicted in panel (g). Statistical test employed: Mann-Whitney test (GraphPad Prism 8). Unless otherwise stated, within the graphs as a distinct P value (if statistically insignificant), the significance intervals are denoted: P value < 0.0001 (****), 0.0001 to 0.001 (***), 0.001 to 0.01 (**) and 0.01 to 0.05 (*). Data relating to each individual embryo assayed are detailed in supplementary tables S1.

**Figure 3 (colour).**
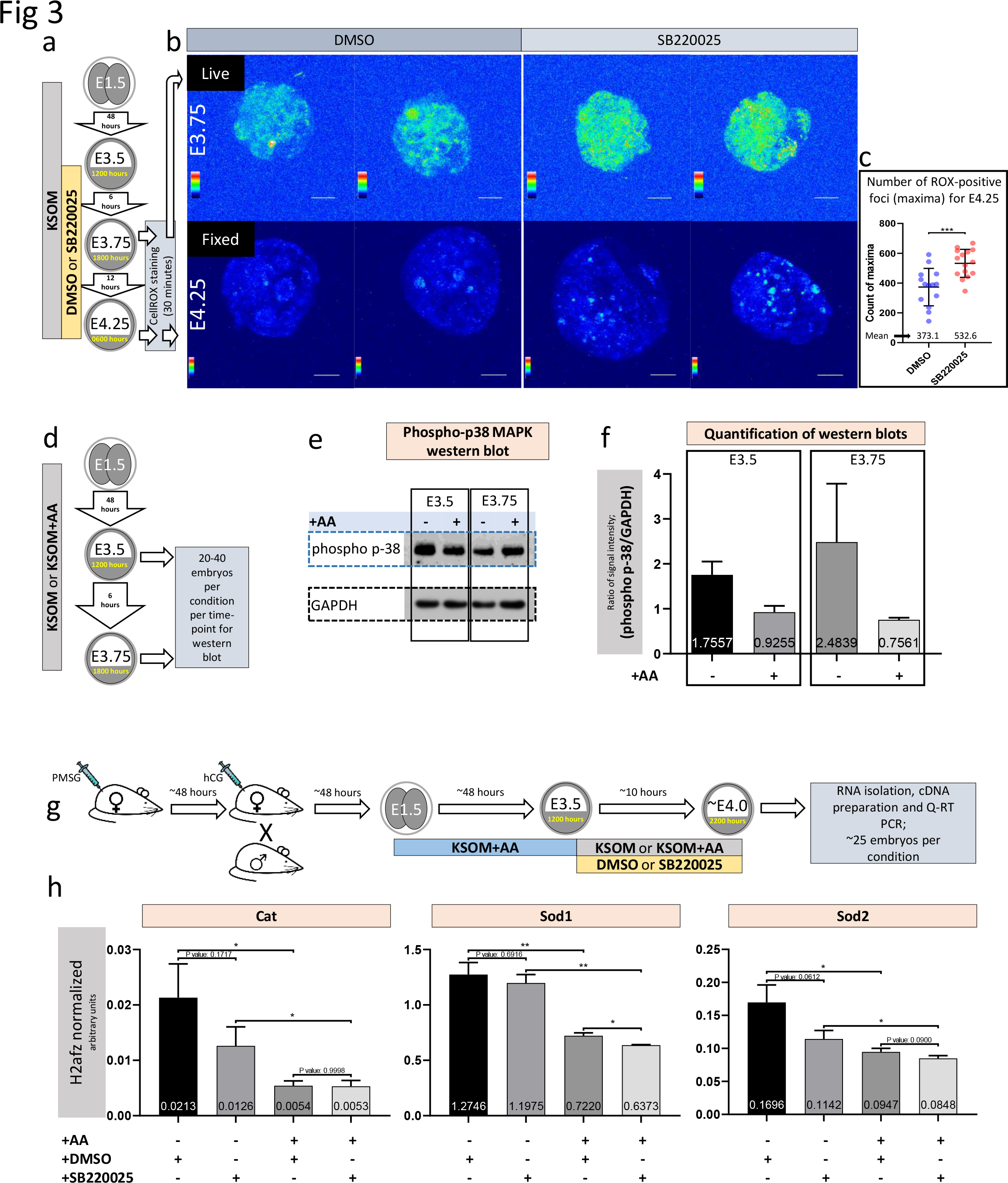
Amino acid starvation coupled with p38-MAPK inhibition induces increased blastocyst ROS levels whereas amino acid starvation alone is sufficient to induce increased phosphorylation/activation of p38-MAPK and transcription of genes with enzymatic anti-oxidant properties. **a.** Experimental design detailing culture conditions for embryos and sampling time points used to visualise and quantify blastocyst ROS levels. **b.** Projected confocal z-stack images of blastocyst embryos stained with CellROX Green at E3.75 and 4.25 and were respectively imaged live or after fixation. A spectral rainbow palette is used to denote signal intensity levels (blue representing lowest to white denoting highest signal intensity); scale bar = 20μm. **c.** Quantification of the average per embryo incidence of ROX-positive foci (maxima) for the E4.25 assayed and fixed embryo groups cultured in KSOM plus DMSO and SB220025; as described in panels a) and b). The statistical test employed was an unpaired, two-tailed students t-test. The stated significance intervals are depicted as: P value < 0.0001 (****), 0.0001 to 0.001 (***), 0.001 to 0.01 (**) and 0.01 to 0.05 (*). The graphs represent dot plots of total sample size together with stated experimental group means and the standard deviations (error bars). **d.** Experimental design detailing culture conditions for embryos and sampling time points used in western blotting based assay of activated and phosphorylated p38-MAPK protein expression in blastocyst embryos cultured in KSOM ±AA. **e.** Representative western blot of phosphorylated p38-MAPK and GAPDH (as control) blastocyst protein levels, after culture in KSOM and KSOM+AA, at E3.5 and E3.75. Note, uneven sample loading (GAPDH) and hence need for normalised quantitation (panel f). **f.** Relative signal intensity quantification data of the GAPDH normalised levels of phosphorylated p38-MAPK protein expression (judged by western blot) in blastocysts (at E3.5 and E3.75) after culture in KSOM and KSOM+AA (from two independent biological replicates). **g.** Experimental design detailing culture conditions for embryos used in Q-RTPCR based quantification of enzymatic anti-oxidant gene mRNA expression in blastocysts (at E4.0) cultured in KSOM±AA treated with DMSO or p38-MAPK inhibitor (SB220025). **h.** Quantification of the relative transcript expression levels of *Cat*, *Sod1* and *Sod2*, internally normalised to *H2afz* levels, across the four conditions. The statistical test employed were Welch’s ANOVA tests followed by Dunnett’s T3 multiple comparisons test (GraphPad Prism 8). Unless otherwise stated within individual charts as a specific P value (if statistically insignificant), the stated significance intervals are as follows: P value < 0.0001 (****), 0.0001 to 0.001 (***), 0.001 to 0.01 (**) and 0.01 to 0.05 (*); error bars denote standard deviation and mean *H2afz* normalised expression levels are numerically stated.

**Figure 4 (colour).**
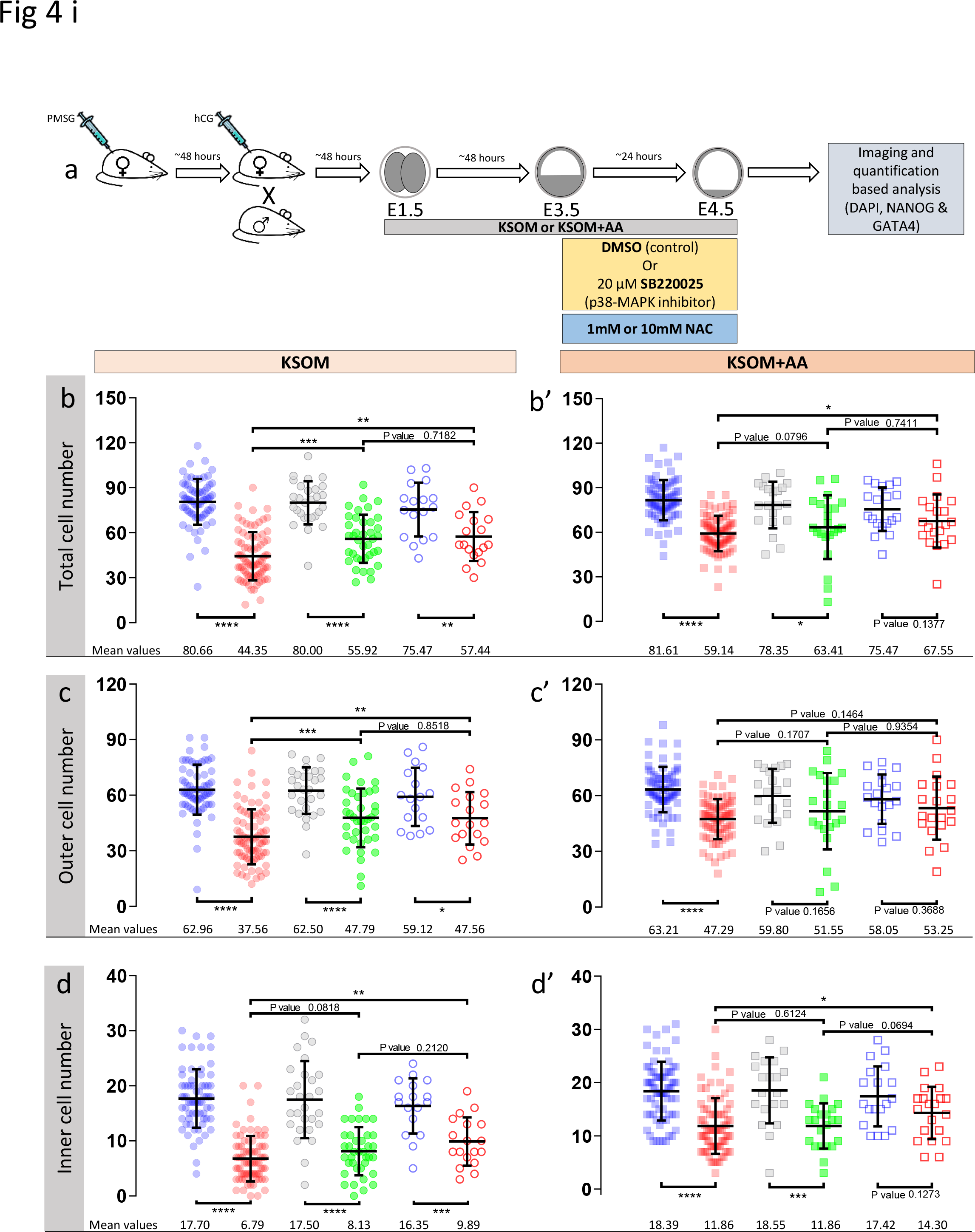
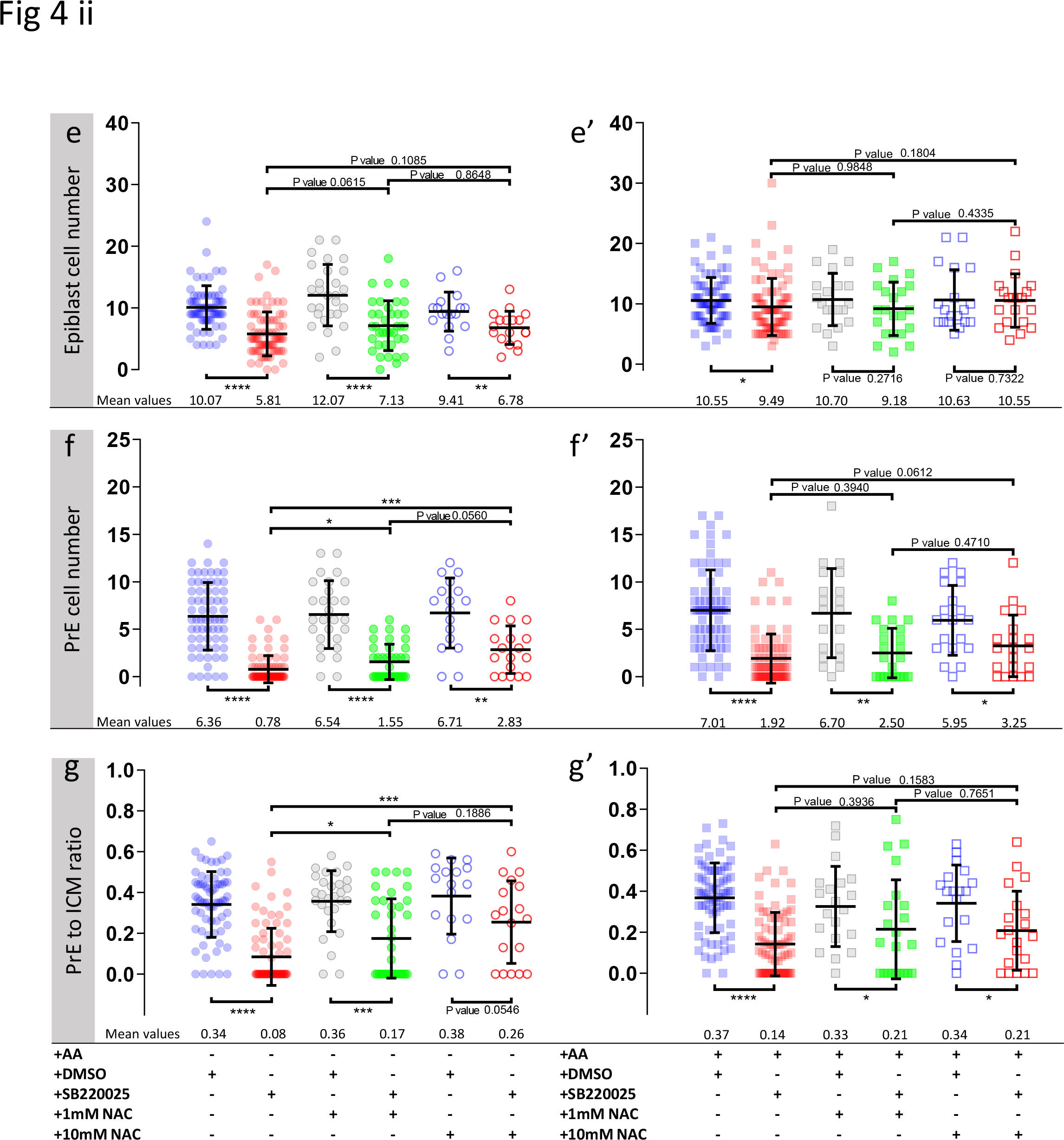
Effect of anti-oxidant (NAC) treatment on p38-MAPK inhibited blastocyst cell numbers, based on DAPI, NANOG and GATA4 staining and confocal microscopy, in culture conditions ± exogenous amino acid supplementation. **a.** Experimental design detailing culture conditions of embryos recovered at the 2-cell (E1.5) stage, in regard to culture media amino acid supplementation status and the possible combined regimes of NAC plus DMSO or p38-MAPK inhibitor (SB220025) treatment from E3.5-E4.5. KSOM group (b-g), left to right: DMSO (n = 67), p38-MAPK inhibition (n = 77), DMSO + 1mM NAC (n = 28), p38-MAPK inhibition + 1mM NAC (n = 38), DMSO + 10mM NAC (n = 17) and p38-MAPK inhibition + 10mM NAC (n = 18). KSOM+AA group (b’-g’), left to right: DMSO (n = 71), p38-MAPK inhibition (n = 76), DMSO + 1mM NAC (n = 20), p38-MAPK inhibition + 1mM NAC (n = 22), DMSO + 10mM NAC (n = 19) and p38-MAPK inhibition + 10mM NAC (n = 20). An explanatory matrix of the composition of each experimental condition is present beneath the relevant charts, at the base of the whole figure. **b & b’.** Quantification of total cell number based on counting DAPI stained nuclei of the blastomeres. **c & c’.** Quantification of outer cell number based on subtracting NANOG and/or GATA4 positive inner cells from the DAPI stained total cell number (plus position within the embryo). **d & d’.** Quantification of inner cell number based on NANOG and/or GATA4 stained inner cells number (plus position within the embryo). **e & e’.** Quantification of epiblast cell number based on NANOG alone stained inner cells. **f & f’.** Quantification of primitive endoderm cell number based on GATA4 alone stained inner cells. **g & g’.** Quantification of the contribution of GATA4 stained primitive endoderm cells as a ratio of the total inner cells (NANOG and/or GATA4 stained cells). Statistical test employed: Mann-Whitney test (GraphPad Prism 8). Unless otherwise stated, within the graphs as a distinct P value (if statistically insignificant), the significance intervals are as follows: P value < 0.0001 (****), 0.0001 to 0.001 (***), 0.001 to 0.01 (**) and 0.01 to 0.05 (*). Note, data relating to DMSO and p38-MAPK inhibition conditions in KSOM and KSOM+AA media are reproduced from Fig. 2 to aid comparison with NAC treated groups. Data relating to each individual embryo assayed are detailed in supplementary tables S2. A Welch’s ANOVA statistical analysis of the data presented in this figure is presented in supplementary table 5.

**Figure 5 (colour).**
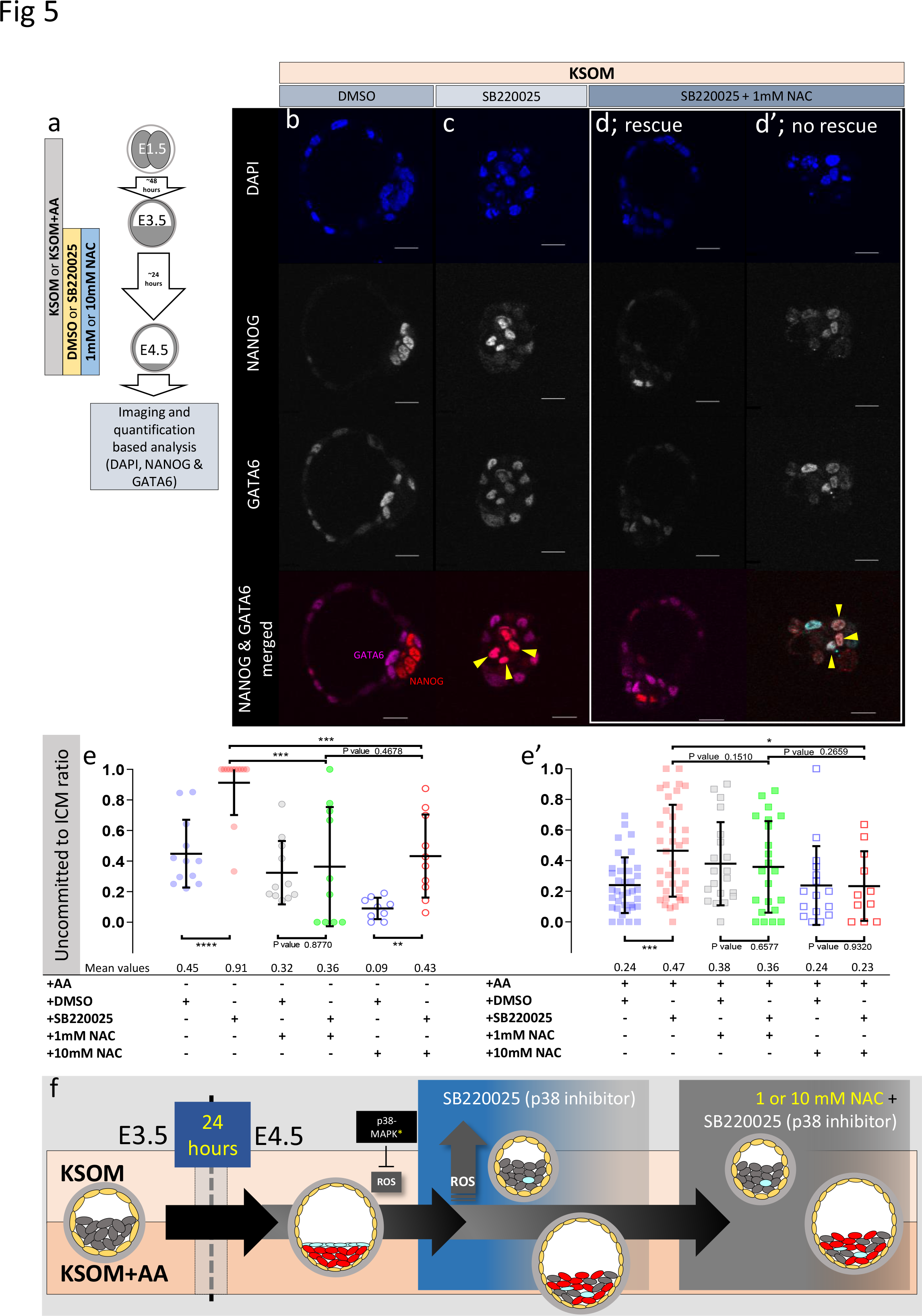
Effect of anti-oxidant (NAC) treatment on uncommitted ICM cells in p38-MAPK inhibited blastocysts, based on DAPI, NANOG and GATA6 staining and confocal microscopy, in culture conditions ± exogenous amino acid supplementation. **a.** Experimental design detailing culture conditions of embryos recovered at the 2-cell (E1.5) stage, in regard to culture media amino acid supplementation status and the possible combined regimes of NAC plus DMSO or p38-MAPK inhibitor (SB220025) treatment from E3.5-E4.5. **b to d’.** Single confocal microscopy z-sections of representative E4.5 embryos, cultured in KSOM (plus indicated combined regimes of DMSO/SB220025 and NAC from E3.5-E4.5) stained for, from top to bottom, nucleus/ DNA (DAPI), epiblast (NANOG), primitive endoderm (GATA6) and uncommitted cells (NANOG and GATA6 co-staining – see pseudo-coloured merged image). Yellow arrowheads highlight ICM cells co-expressing both markers (*i.e.* uncommitted cells); panels (c) and (c’) show examples of embryos in which addition of 1mM NAC was able to either rescue, or not rescue, ICM cells from the p38-MAPK inhibition induced uncommitted cell fate, respectively. All images in one vertical panel are of the same embryo at same magnification. Scale bar = 20μm. **e & e’.** Contribution of GATA6 and NANOG co-stained (*i.e.* uncommitted) cells, shown as a ratio of the averaged total of inner cells (NANOG and/or GATA6 inner stained cells), in embryos cultured in KSOM (e) or KSOM+AA (e’) under indicated (see explanatory matrix, as in Fig. 4) combined regimes of DMSO/SB220025 and NAC treatment from E3.5-E4.5; KSOM (e), left to right: DMSO (n = 12), p38-MAPK inhibition (n = 12), DMSO + 1mM NAC (n = 11), p38-MAPK inhibition + 1mM NAC (n = 10), DMSO + 10mM NAC (n = 9) and p38-MAPK inhibition + 10mM NAC (n = 10); KSOM+AA (e’), left to right: DMSO (n = 37), p38-MAPK inhibition (n = 38), DMSO + 1mM NAC (n = 20), p38-MAPK inhibition + 1mM NAC (n = 22), DMSO + 10mM NAC (n = 16) and p38-MAPK inhibition + 10mM NAC (n = 11). Statistical test employed: Mann-Whitney test (GraphPad Prism 8). Unless otherwise stated, within the graphs as a distinct P value (if statistically insignificant), the significance intervals are: P value < 0.0001 (****), 0.0001 to 0.001 (***), 0.001 to 0.01 (**) and 0.01 to 0.05 (*). Data relating to each individual embryo assayed are detailed in supplementary table S3. A Welch’s ANOVA statistical analysis of the data presented in this figure is presented in supplementary table 5. **f.** Model: irrespective of exogenous AA media supplementation (*i.e.* KSOM±AA), early (E3.5) stage blastocysts are able to appropriately specify and segregate ICM cell lineages (red EPI & blue PrE) by the late blastocyst (E4.5) stage. However, inhibition of p38-MAPK reveals derivation of ICM lineages from initially uncommitted progenitors (shown in grey) is differentially impaired in a manner dependent on exogenous AA media supplementation. Whereas in the presence of exogenous AA (KSOM+AA) it is only the PrE (and not EPI) that fails to specify, a complete absence of provided AA (KSOM) is associated with the majority of ICM cells (*i.e.* EPI & PrE progenitors) being retained in the uncommitted state by the late blastocyst (E4.5) stage, with reduced cell number and smaller cavities. Further supplementation of the anti-oxidant NAC is able to partially rescue the more severe phenotype associated with p38-MAPK inhibition in a subset of embryos cultured in KSOM media (to a point resembling p38-MAPK inhibition in KSOM+AA media; *i.e.* specified EPI, unspecified PrE), whereas NAC addition had no effect on the milder p38-MPAK inhibition phenotype observed in embryos cultured in KSOM+AA. Collectively, such data indicate a requirement for p38-MAPK to homeostatically buffer, AA depletion induced oxidative stress (caused by increased ROS levels), to allow germane blastocyst development and conditions conducive to EPI specification; however PrE specification and ultimate differentiation is governed by an, as yet unknown, but independent p38-MAPK mediated mechanism [downstream of FGF-signalling, as previously described (Thamodaran and Bruce, 2016)].

### Quantitative real-time PCR (Q-RTPCR)

2-cell stage (E1.5) embryos were collected and cultured in KSOM+AA until E3.5 (*i.e.* at 12:00 hours) stage and then equally distributed in to four experimental pre-equilibrated media conditions (*i.e.* i. KSOM +DMSO, ii. KSOM +SB220025, iii. KSOM+AA +DMSO and iv. KSOM+AA +SB220025). At 22:00 hours (*i.e.* 10 hours of treatment), 25 embryos from each condition were collected immediately processed for RNA extraction and isolation using the ARCTURUS® PicoPure® RNA Isolation Kit (Applied Biosystems™; catalogue number KIT0204), following the manufacturer’s protocol. The entire eluted volume of total RNA was immediately DNase treated with Invitrogen™ TURBO™ DNase (Catalogue number: AM2238) according to the manufacturer provided protocol. The whole sample was then subject to cDNA synthesis using Invitrogen™ SuperScript™ III Reverse Transcriptase (Catalogue number: 18080044), as directed by the manufacturer and employing Invitrogen™ Oligo d(T)16 (Catalogue number: N8080128), ThermoScientific™ dNTP Mix (Catalogue number: R0192) and Applied Biosystems™ RNase Inhibitor (Catalogue number: N8080119). The final cDNA volume of 30μl was diluted to 45μl with nuclease free water and 1μl used in 10μl individual final SYBR-green based Q-RT-PCR reaction volumes (qPCRBIO SyGreen Mix Lo-ROX - Catalogue number: PB20.11). A Bio-Rad CFX96 Touch Real-Time PCR Detection System apparatus, employing standard settings, was employed for data accumulation and initial analysis was performed with the accompanying Bio-Rad CFX Manager™ software. Triplicate measurements per gene (the sequence of individual oligonucleotide primers, used at a final concentration of 300nM, are provided in supplementary table S6) were assayed from two biological replicates that were each technically replicated. The averaged transcript levels of analysed genes (*i.e. Cat*, *Sod1* and *Sod2*) were derived after internal normalisation against *H2afz* mRNA levels, in four experimental culture conditions assayed. As such data was acquired and initially analysed with CFX Manager™, then processed in Microsoft Excel (biological and technical replicate averaging) and GraphPad Prism 8 (graphical output). Welch’s ANOVA statistical significance test followed by Dunnett’s T3 multiple comparisons test were employed. Unless otherwise stated within individual graphs as a specific P value (if statistically insignificant), the stated significance intervals were depicted as: P value < 0.0001 (****), 0.0001 to 0.001 (***), 0.001 to 0.01 (**) and 0.01 to 0.05 (*); error bars denote calculated standard deviations.

### Blastocyst Reactive Oxygen Species (ROS) staining

Collected 2-cell stage embryos were cultured in KSOM until E3.5 (12:00) then moved to either KSOM +DMSO or KSOM +SB220025 (pre-equilibrated for 3 hours prior) and cultured for another 6 hours (*i.e.* E3.75; alternatively expressed as 18:00 hours on the same day) or to E4.25 (*i.e.* until 06:00 hours the next day). Thereafter embryos were transferred to M2 media containing 5μM Invitrogen™ CellROX™ Green Reagent (a ROS specific reporter dye; catalogue number: C10444), that had been pre-equilibrated in the dark at 37°C for 30 minutes, and incubated under the same conditions for 30 minutes before being washed through two drops of regular M2 media. Whereas E3.75 (n=2 per experimental group) embryos were immediately live mounted M2 drops and imaged under the confocal microscope (FV10i, Olympus®, using appropriate preparatory CellROX™ Green filter settings), those ROS stained embryos collected at E4.25 (KSOM +DMSO, n=14; KSOM +SB220025, n=15) were fixed in 4% paraformaldehyde prior to confocal microscopic imaging. Confocal images are depicted as projections of individual z-stack images using the FV10i-SW image acquisition software (Olympus®) rainbow spectral intensity palette (from blue to white, representing lowest to highest signal intensities). All the images in each group (*i.e.* E3.75 and E4.25) were acquired at equal laser intensity and detector sensitivity. The number of ROX-positive/staining foci in the z-stack projections of individual embryos fixed at E4.25 were quantified using Image J (http://rsbweb.nih.gov/ij/), by first subtracting background (using a rolling ball radius of 50 pixels) and invoking the ‘finding maxima tool’ (prominence>5). The counted number of foci were statistically verified and graphically depicted using GraphPad Prism 8; statistical test employed was an unpaired, two-tailed t-tests. Unless otherwise stated within individual graphs as a specific P value (if statistically insignificant), the stated significance intervals are depicted: P value < 0.0001 (****), 0.0001 to 0.001 (***), 0.001 to 0.01 (**) and 0.01 to 0.05 (*). The graphs represent dot plot of total sample size together with mean and the standard deviation bars indicated.

### Phosphorylated/ activated p38-MAPK western blotting and quantification

Collected 2-cell stage embryos were cultured in either KSOM or KSOM+AA until E3.5 (12:00 hours) or E3.75 (18.00 hours) and 20-40 embryos processed for western/immunoblotting per culture condition; briefly embryos were washed through two drops of Dulbecco’s PBS (Sigma-Aldrich; catalogue number: BSS-1005-B), transferred to 1.5 ml microfuge tubes (removing excess PBS) and flash frozen in liquid nitrogen. To prepared samples 10 μL of 10× SDS reducing agent/loading buffer (NuPAGE buffer, ThermoFisher Scientific, NP 0004, ThermoFisher Scientific) was added and then boiled at 100°C for 5 minutes. Loaded proteins were then electrophoretically separated on gradient precast 4–12% SDS–PAGE gels (ThermoFisher Scientific, NP0323) and transferred to Immobilon P membranes (Merck group, IVPD00010) using a semi-dry blotting system (Biometra/ Analytik Jena) for 25 minutes at 5mA/ cm^2^. Blotted membranes were blocked in 5% skimmed milk powder dissolved in 0.05% Tween-Tris pH 7.4 buffered saline (TTBS), for 1 hour, briefly rinsed in TTBS and then incubated overnight at 4°C overnight in 1% milk/TTBS containing primary antibody (against phosphorylated p38-MAPK). Membranes were washed in three changes of TTBS buffer (20 minutes each at room temperature) and anti-immunoglobulin-species-specific-peroxidase conjugated secondary antibody added to the blot in 1% milk/TTBS, for 1 hour (room temperature). Immuno-detected proteins were visualized by chemiluminescent photographic film exposure (ECL kit; GE Life Sciences, RPN2232) and digitally scanned using a GS-800 calibrated densitometer (Bio-Rad Laboratories) and quantified using ImageJ (http://rsbweb.nih.gov/ij/). Antibody stripped membrane blots were re-probed and quantified, for loading controls (detecting GADPH), in an identical manner. Note, presented quantified data (Fig. 3f) of GAPDH normalised phosphorylated p38-MAPK levels are taken from two independent biological replicates. Supplementary table S4 details the identity and utilised concentrations of the primary and peroxidase-conjugated antibodies used.

## Results & Discussion

### p38-MAPK activity buffers amino acid availability to ensure germane blastocyst maturation and appropriate ICM cell lineage derivation

As outlined above, we assayed the effect of exogenous AA supplementation on *in vitro* mouse blastocyst formation, assaying total, outer/TE, overall ICM, pluripotent EPI and PrE cell numbers. Concomitantly, we analysed the effect of pharmacological p38-MAPK inhibition [using SB220025 (Jackson et al., 1998)], given our previously identified role for p38-MAPK in regulating PrE specification and formation during the blastocyst maturation developmental window [E3.5-E4.5 (Thamodaran and Bruce, 2016)]. Accordingly, 2-cell stage mouse embryos were *in vitro* cultured until the early blastocyst stage (E3.5) in a commonly utilised and chemically defined commercial growth media lacking AAs (except L-glutamine; KSOM) or the identical media supplemented with essential and non-essential AAs (KSOM+AA). Embryos were then switched to the equivalent media containing either SB220025 or DMSO (vehicle control), further cultured to the late blastocyst stage (E4.5), fixed and immuno-fluorescently stained for EPI (NANOG) or PrE (GATA4) marker protein expression. Irrespective of AA supplementation status, we did not observe any morphological differences between the DMSO control treated groups; each yielding hatching blastocysts with appropriately large cavities (Fig. 1b and b’) and ICMs consisting a NANOG positive (NANOG+) EPI compartment overlaid with mono-layered GATA4 positive (GATA4+) PrE cells (Fig. 1d and f). Neither the average number of total, outer and inner (including PrE and EPI) cells, nor the PrE:total ICM cell number ratio, were significantly different between each of the DMSO treated control groups (Fig. 2). Thus, exogenous AA supplementation did not overtly affect embryonic/blastocyst development or ICM lineage derivation during the preimplantation period (notwithstanding possible undetectable changes, possibly epigenetic in origin, that may potentially underpin any subsequent DOHaD phenotypes). Similar inspection of the p38-MAPK inhibited groups confirmed our previously reported data (Thamodaran and Bruce, 2016), whereby blastocysts were morphologically smaller (particularly the KSOM group) (Fig. 1c and c’) and had a robust PrE deficit (note lack of GATA4+ cells Fig. 1e, e’, g and g’). Indeed, a detailed analysis of cell numbers (Fig. 2) revealed the p38-MAPK inhibited embryos that were cultured in KSOM were significantly more adversely affected than those cultured in KSOM+AA, comprising fewer overall, outer and inner cells, although both inhibited groups had significantly fewer cells than their equivalent DMSO controls. Furthermore, p38-MAPK inhibited embryos cultured in KSOM also had significantly fewer EPI cells (5.81 cells) than either p38-MAPK inhibited blastocysts cultured in KSOM+AA (9.49 cells) or the appropriate DMSO treated KSOM control groups (10.07 cells) (Fig. 2d). This trend was also observed in the PrE, albeit representing a small difference in the overall magnitude; *i.e.* an average difference of 1.14 cells between the KSOM+AA and KSOM conditions under p38-MAPK inhibition (0.78 cells in KSOM and 1.92 in KSOM+AA, compared with 6.36 and 7.01 in the respective control DMSO conditions - Fig. 2e). p38-MAPK inhibition, thus, severely attenuated PrE differentiation irrespective of AA supplementation status, with the magnitude of the effect being marginally greater, yet reaching statistical significance, in blastocysts matured in KSOM media. Interestingly, there was no similar robust reduction in EPI cells in p38-MAPK inhibited embryos from the KSOM+AA cultured group (10.55 cells in control vs. 9.49 in inhibited conditions), in line with our previous observations (Thamodaran and Bruce, 2016). Collectively, these data confirm maximally reduced cell number phenotypes associated with p38-MAPK inhibition under non-supplemented KSOM culture conditions, and indicate a developmental buffering capacity of active p38-MAPK (that potentially ensures required AA availability) that is necessary for appropriate blastocyst development/maturation. However, the fact that exogenous AA supplementation is able to elicit a near complete rescue EPI cell number deficits caused by p38-MAPK inhibition but only has a marginal effect on robustly impaired PrE differentiation, demonstrates such a regulative AA-related homeostatic role of p38-MAPK to be distinct and independent of that it fulfils in potentiating PrE differentiation (Thamodaran and Bruce, 2016).

How active p38-MAPK executes this homeostatic role in the absence of exogenous AAs (see KSOM +DMSO conditions Figs.1&2) is unclear but may involve sequestration of intra-cellular sources of AA via regulated autophagy, as reported in other non-embryo-related models/systems (Corcelle et al., 2007;Webber, 2010;Webber and Tooze, 2010;Henson et al., 2014). Consistently, p38-MAPK has also been reported to regulate mTOR containing complexes, involved in balanced metabolism, cell growth/proliferation control and autophagy (Casas-Terradellas et al., 2008;Cully et al., 2010;Wu et al., 2011;Gutierrez-Uzquiza et al., 2012;Linares et al., 2015). Moreover, partial mTOR inhibition in mouse blastocysts is known to induce a state of developmental diapause (Bulut-Karslioglu et al., 2016), whilst relative differences in mTOR activity (linked to p53 activity) have been reported as a mechanism by which cells exiting the naive pluripotent state compete and are potentially eliminated from early post-implantation embryonic tissues (Bowling et al., 2018). It would be interesting to investigate further the potential regulation of mTOR via p38-MAPK. Interestingly, there also exists precedent from cell line models for atypical glucose induced autophagy, which is independent of mTOR and relies on p38-MAPK. This mechanism of glucose induced and p38-MAPK dependant autophagy is however only operative under conditions of nutrient deprivation, such as depletion of exogenously provided AAs (Moruno-Manchon et al., 2013). Given the base media (*i.e.* KSOM) used in this study contains glucose, it is possible a similar mechanism of induced and p38-MAPK dependent autophagy is responsible for the overtly normal blastocyst maturation observed in DMSO treated control embryos cultured in the AA-free non-supplemented KSOM.

### p38-MAPK counteracts amino acid depletion induced oxidative stress during blastocyst maturation

AA starvation is closely linked to increased oxidative stress (Harding et al., 2003), whereby induced anti-oxidant mechanisms, involving *de novo* protein expression, are impaired (Vucetic et al., 2017). Additionally, activated p38-MAPK (specifically p38α) has been reported to orchestrate the induced/stabilised expression of enzymatic antioxidants (Gutierrez-Uzquiza et al., 2012). Therefore, given the morula to blastocyst transition in mouse preimplantation development is accompanied by a large increase in glucose utilisation and oxygen consumption (Brown and Whittingham, 1991;Leese, 2012), with the potential to contribute elevated levels of reactive oxygen species (Murphy, 2009;Harvey, 2019), we hypothesised the observed aggravated effect of p38-MAPK inhibition in the absence of AA supplementation was contributed by increased oxidative stress. In support of this model, we could directly detect increased levels of ROS (using a ROX-dye) in blastocysts under p38-MAPK inhibited conditions in both live E3.75 –(Fig. 3b, upper panels) and fixed E4.25 (Fig. 3b, lower panels and Fig. 3c, detailing quantification of the average number of ROX-positive foci per embryo) stage embryos, cultured in non-supplemented KSOM (see also supplementary Fig. S1, detailing projected z-section confocal micrographs of all ROX stained E4.25 stage fixed embryos used in ROS quantification). Additionally, a comparative and quantitative assay of the levels of functionally active phosphorylated p38-MAPK protein in blastocysts derived after culture in either KSOM or KSOM+AA, reported enhanced levels of active p38-MAPK in the KSOM cultured group condition at both E3.5 and E3.75 (Fig. 3e and f); *i.e.* the culture condition, with AA supplementation, already shown to be most sensitive to the effects of p38-MAPK inhibition (Figs. 1 and 2). In further support of the model, we could also detect significantly enhanced expression of recognised mRNA transcripts for enzymatic antioxidants [*i.e.* Catalase/ *Cat* and Superoxide-dismutases 1 & 2/ *Sod1* & *Sod2* (Gutierrez-Uzquiza et al., 2012)] in control DMSO treated blastocysts cultured in un-supplemented KSOM versus KSOM+AA. Moreover, in the case of *Cat* and *Sod2* expression, such comparatively enhanced levels could be attenuated by p38-MAPK inhibition, although not to levels observed in blastocysts cultured in KSOM+AA (Fig. 3c – note all blastocysts were initially cultured in KSOM+AA until E3.5, before being transferred to one of the four assayed media conditions). Collectively, these data support the hypothesis that increased oxidative stress caused by a lack of exogenous AA media supplementation is buffered by active p38-MAPK to ensure germane blastocyst development and the potential to adopt appropriate ICM cell fates.

Accordingly, we therefore tested if blastocyst cell number deficits caused by p38-MAPK inhibition could be ameliorated by providing exogenous anti-oxidants to the culture media. Thus, we repeated the cell counting experiments described above, including extra conditions in which cultured blastocysts (E3.5-E4.5), under control (DMSO) and p38-MAPK inhibited conditions, were provided the antioxidant N-acetyl-cysteine (NAC; either 1mM or 10mM - Fig. 4). Focussing on the non-supplemented (KSOM) group, we observed that NAC addition to control DMSO treated embryo groups had no significant effect on average total, outer or inner cell number; nor EPI/ PrE derivation (*n.b.* PrE:ICM ratio; Fig. 4b-g and supplementary table S5: Welch’s-ANOVA test), suggesting a lack of significant oxidative stress under control conditions. However, under p38-MAPK inhibited conditions, NAC supplementation caused significant increases (*i.e.* ‘rescues’) in all but EPI cell numbers (Fig. 4b-g and supplementary table S5), when compared with p38-MAPK inhibition in the absence of NAC; indicating a role for p38-MAPK in mitigating depleted AA induced oxidative stress. A marked divergent effect was indeed observed in respect to the two ICM cell lineages. Whilst, NAC treatment did not significantly alter the average number of EPI (NANOG+) cells under p38-MAPK inhibition (5.81, 7.13 and 6.78 in non-supplemented, 1mM and 10mM NAC conditions respectively; Fig. 4e), the number of PrE cells (GATA4+) was significantly increased (0.78, 1.55 and 2.83 in non-supplemented, 1mM and 10mM NAC conditions respectively; Fig. 4f), with an improving trend towards higher concentrations of NAC (Fig. 4f and g). Therefore, these data strongly imply that under conditions of exogenous AA depletion, p38-MAPK activity specifically supports the development of differentiating extraembryonic lineages (TE and PrE) by combating induced oxidative stress (as revealed by the described rescue phenotypes associated with concomitant p38-MAPK inhibition and NAC supplementation). Moreover, the data also suggest this anti-oxidant role is not extended to the pluripotent EPI and confirm that PrE differentiation, *per se*, is sensitive to oxidative stress. These data accord with our previous observations that p38-MAPK inhibition specifically impairs PrE cell-fate derivation, by preventing resolution of initially uncommitted cell fate states within early blastocyst ICMs, without significantly influencing EPI formation (Thamodaran and Bruce, 2016) (recapitulated here; KSOM+AA - Figs.2 & 4). However, the observed NAC-derived rescue effects were only partial when compared to average cell numbers recorded in the appropriate DMSO control groups, and with the exception of GATA4+ PrE derivation, were not further enhanced by the higher (10mM) NAC concentration. Indeed, increased/rescued total, outer and PrE cell numbers in NAC treated p38-MAPK inhibited embryos, in the KSOM group, were only equivalent to those observed in the corresponding KSOM+AA conditions (Fig. 4). This suggests further, as yet unknown, p38-MAPK regulated mechanisms (unrelated to oxidative stress) must also contribute to ensure appropriate numbers of total, outer (TE) and PrE cells during blastocyst maturation. However, the present data confirm the existence of at least a dual role of p38-MAPK during blastocyst maturation. Firstly, that concerned with homeostatic regulation of AA availability and associated oxidative stress and secondly, that by which p38-MAPK directly regulates PrE specification/ differentiation, as previously described (Thamodaran and Bruce, 2016).

It is noteworthy that a study employing cancer cell line models reports p38-MAPK as elevating glucose uptake but shunting its metabolism away from glycolysis in favour of the pentose phosphate pathway to generate increased levels of NADPH that are required to counteract reactive oxygen species (ROS) (Desideri et al., 2014). Thus, it is tempting to speculate a similar mechanism to counteract ROS, under depleted AA conditions, may also be operative in the blastocyst. Interestingly, recently available preprint study data compellingly report the importance of blastocyst cavity expansion for PrE specification (https://doi.org/10.1101/575282); given such expansion requires a functioning TE (Madan et al., 2007), it is possible any impairment in TE cell number/ function could negatively impact this process. Consistently, we have observed the blastocyst cavities of p38-MAPK inhibited blastocysts to be visibly smaller than controls (Fig. 1b to c’; highlighted by black arrowheads), irrespective of AA supplementation status. Reports relating to glioblastoma cancer models have also clearly uncovered p38-MAPK dependant mechanisms of cell survival, involving induction of the *Cox2* gene (catalysing the committed step in prostaglandin synthesis) (Parente et al., 2013) and that have been linked to AA starvation (Li et al., 2017). However, it is improbable p38-MAPK exerts a similar role in the mouse blastocyst ICM, as we were unable to detect any basal or induced, as the recognised mechanism of regulated expression (Smith et al., 2000), *Cox2* derived mRNA transcripts in either the control or experimental conditions (as stated in Fig. 3h), again irrespective of KSOM media AA supplementation status. Lastly, we found the addition of NAC to p38-MAPK inhibited blastocysts under KSOM+AA culture conditions had no significant effects (Fig. 4b’-f’ and supplementary table S5), suggesting AA supplementation alone is sufficient to mitigate deleterious oxidative stresses (potentially caused by increased ROS production) associated with p38-MAPK inhibition. However, the fact p38-MAPK inhibition was still associated with significantly reduced total, outer and PrE cell numbers further supports the existence of additional uncharacterised, yet p38-MAPK mediated, mechanism(s) of supporting appropriate blastocyst cell number.

### p38-MAPK dependent homeostatic regulation of amino acid availability, and associated depletion induced oxidative stress, facilitates PrE progenitor specification from an initially uncommitted state of cell fate, during blastocyst maturation

Cells of maturing mouse blastocysts ICM (E3.5 - E4.5) transit from an initially uncommitted state, expressing both NANOG (EPI) and GATA6 (PrE), via an intermediate stage whereby progenitors of each lineage express either marker in a randomly distributed and mutually exclusive manner (the so-called ‘salt & pepper’ pattern), culminating in the separated late blastocyst tissue layers (PrE having initiated further marker protein expression; *e.g.* SOX17 and GATA4 (Kuo et al., 1997;Chazaud et al., 2006;Niakan et al., 2010)). We previously reported p38-MAPK inhibition during this developmental window significantly impairs PrE (GATA4+), but not EPI (NANOG+), formation by retaining PrE progenitor cells in the uncommitted state (Thamodaran and Bruce, 2016). Given such experiments were performed in KSOM+AA, we now decided to also assay PrE specification from the uncommitted state in KSOM cultured blastocysts, in the presence/absence of NAC. Consistent with our previous study (Thamodaran and Bruce, 2016), the inhibition of p38-MAPK in blastocysts cultured in KSOM+AA was associated with increased numbers of uncommitted/ PrE progenitor ICM cells (co-expressing NANOG and GATA6 - illustrated by an increase in the uncommitted cell:total ICM ratio– Fig. 5e’). However, NAC supplementation resolved this uncommitted state to levels observed in corresponding DMSO treated controls (reaching significance at 10mM NAC). These data suggest p38-MAPK inhibition in KSOM+AA media is associated with levels of oxidative stress, that represent an impediment to PrE progenitor specification in the blastocyst (*i.e. a* resolution of the uncommitted state by downregulating NANOG and maintaining GATA6 expression). Moreover, they invoke an endogenous anti-oxidant role for p38-MAPK that promotes PrE specification, although crucially such a role is not alone sufficient to ensure full differentiation as marked by GATA4 expression (discussed above - Fig. 4). Whereas in KSOM cultured p38-MAPK inhibited blastocysts, the observed uncommitted ICM phenotype (NANOG and GATA6 co-expression) was much more robust (affecting virtually all ICM cells - *i.e.* progenitors of PrE and EPI alike; Fig. 5c & e). Such data suggest a lack of AA supplementation itself, coupled with p38-MAPK inhibition, severely impairs resolution of the uncommitted state towards either PrE or EPI (even if some very limited initiation of GATA4 expression can occur later - Figs.1, 2 & 4). This is opposed to the effect of p38-MAPK inhibition in KSOM+AA, whereby only resolution of uncommitted cells to PrE is impaired [Fig. 5e’ (Thamodaran and Bruce, 2016)]. Addition of NAC to DMSO treated control KSOM culture groups revealed that the antioxidant alone, in the absence of supplemented AAs, promotes resolution of the uncommitted state (reaching significance at 10mM NAC), although as reference above this is not translated to full PrE differentiation (as marked by GATA4 expression – Figs. 1, 2 & 4). However, the effect is blunted under p38-MAPK inhibited conditions with only a sub-population of embryos resolving NANOG and GATA6 co-expression patterns (Fig. 5d & d’).

Therefore, AA supplementation can ameliorate developmentally persistent, p38-MAPK inhibition associated, uncommitted ICM/PrE cell fates that can be further augmented by additional anti-oxidant treatment using NAC. As NAC mediated resolution of such p38-MAPK inhibition induced uncommitted states (affecting EPI and PrE progenitors) is not as efficient in the absence of exogenous AA, a PrE specification/ differentiation promoting role of p38-MAPK, regulating appropriate amino acid homeostasis and counteracting oxidative stress, is further supported.

## Conclusions

Our data reveal novel AA sensing and regulative/ homeostatic buffering mechanisms, centred on p38-MAPK, that counteract oxidative stress and ensure germane mouse blastocyst development; manifest in appropriate cell number and resolution of uncommitted EPI/PrE progenitors within the ICM (summarised in Fig. 5f). These findings resonate with DOHaD models, illustrating how varying nutritional status can place regulatory/metabolic burdens upon the early embryo, with consequences for the differentiation of an extraembryonic tissue, later relied upon during *in utero* development of the foetus. The results also provide important considerations for improved assisted reproductive technologies that necessitate *in vitro* human embryo culture.

## Contributions & acknowledgements

Project contributions; experimental design (P.B., T.V. & A.W.B.), practical research (P.B. & T.V.), western blotting experiments (P.B. & A.Š.), data analysis (P.B. & A.W.B.) and manuscript preparation (P.B. & A.W.B). We acknowledge Institute of Parasitology, Biology Centre (in České Budějovice), Czech Academy of Sciences for housing mice, Marta Gajewska (Institute of Oncology, Warsaw, Poland) and Anna Piliszek (Institute of Genetics & Animal Breeding, Polish Academy of Sciences, Jastrzębiec, Poland) for founder CBA/W mice, Alena Krejčí (Faculty of Science, University of South Bohemia, Czech Republic) for pooling resources and other members of our laboratory (LEMDB – particularly Lenka Gahurová) for valuable inputs and discussions. This work was supported by grants from the Czech Science Foundation/GAČR (18-02891S) and Grant Agency of the University of South Bohemia (GAJU; 012/2019/P).

