## Supplementary data for "p38-mitogen activated kinases mediate a developmental regulatory response to amino acid depletion and associated oxidative stress in mouse blastocyst embryos"

Supplementary table S1

### KSOM

#### DMSO

| # | TOTAL NUMBER OF CELLS | | ICM | | | | | Ratio of PrE/Total ICM | Ratio of Inner/ Outer cells |
|---|---|---|---|---|---|---|---|---|---|
| | Embryo | Double Negative | NANOG +ve | GATA4 +ve | NANOG/ GATA4 double +ve | PrE | Total | | |
| 1 | 55 | 51 | 4 | 0 | 0 | 0 | 4 | 0.00 | 0.08 |
| 2 | 74 | 55 | 10 | 2 | 7 | 2 | 19 | 0.11 | 0.35 |
| 3 | 97 | 77 | 11 | 9 | 0 | 9 | 20 | 0.45 | 0.26 |
| 4 | 85 | 65 | 12 | 7 | 1 | 7 | 20 | 0.35 | 0.31 |
| 5 | 73 | 58 | 12 | 3 | 0 | 3 | 15 | 0.20 | 0.26 |
| 6 | 77 | 57 | 13 | 7 | 0 | 7 | 20 | 0.35 | 0.35 |
| 7 | 86 | 72 | 6 | 5 | 3 | 5 | 14 | 0.36 | 0.19 |
| 8 | 78 | 58 | 9 | 10 | 1 | 10 | 20 | 0.50 | 0.34 |
| 9 | 88 | 59 | 19 | 6 | 4 | 6 | 29 | 0.21 | 0.49 |
| 10 | 96 | 84 | 12 | 0 | 0 | 0 | 12 | 0.00 | 0.14 |
| 11 | 96 | 78 | 8 | 10 | 0 | 10 | 18 | 0.56 | 0.23 |
| 12 | 93 | 79 | 10 | 3 | 1 | 3 | 14 | 0.21 | 0.18 |
| 13 | 81 | 58 | 16 | 6 | 1 | 6 | 23 | 0.26 | 0.40 |
| 14 | 102 | 91 | 7 | 4 | 0 | 4 | 11 | 0.36 | 0.12 |
| 15 | 98 | 77 | 9 | 12 | 0 | 12 | 21 | 0.57 | 0.27 |
| 16 | 72 | 55 | 11 | 4 | 2 | 4 | 17 | 0.24 | 0.31 |
| 17 | 85 | 58 | 13 | 12 | 2 | 12 | 27 | 0.44 | 0.47 |
| 18 | 24 | 9 | 10 | 4 | 1 | 4 | 15 | 0.27 | 1.67 |
| 19 | 87 | 67 | 11 | 6 | 3 | 6 | 20 | 0.30 | 0.30 |
| 20 | 83 | 65 | 9 | 5 | 4 | 5 | 18 | 0.28 | 0.28 |
| 21 | 61 | 45 | 7 | 3 | 6 | 3 | 16 | 0.19 | 0.36 |
| 22 | 87 | 71 | 9 | 6 | 1 | 6 | 16 | 0.38 | 0.23 |
| 23 | 90 | 74 | 9 | 7 | 0 | 7 | 16 | 0.44 | 0.22 |
| 24 | 72 | 55 | 9 | 8 | 0 | 8 | 17 | 0.47 | 0.31 |
| 25 | 79 | 62 | 8 | 8 | 1 | 8 | 17 | 0.47 | 0.27 |
| 26 | 97 | 79 | 9 | 9 | 0 | 9 | 18 | 0.50 | 0.23 |
| 27 | 83 | 61 | 10 | 11 | 1 | 11 | 22 | 0.50 | 0.36 |
| 28 | 70 | 53 | 10 | 6 | 1 | 6 | 17 | 0.35 | 0.32 |
| 29 | 83 | 63 | 9 | 11 | 0 | 11 | 20 | 0.55 | 0.32 |
| 30 | 81 | 63 | 10 | 8 | 0 | 8 | 18 | 0.44 | 0.29 |
| 31 | 63 | 48 | 9 | 5 | 1 | 5 | 15 | 0.33 | 0.31 |
| 32 | 82 | 62 | 11 | 9 | 0 | 9 | 20 | 0.45 | 0.32 |
| 33 | 84 | 70 | 8 | 6 | 0 | 6 | 14 | 0.43 | 0.20 |
| 34 | 77 | 60 | 11 | 6 | 0 | 6 | 17 | 0.35 | 0.28 |
| 35 | 82 | 65 | 6 | 11 | 0 | 11 | 17 | 0.65 | 0.26 |
| 36 | 106 | 86 | 9 | 11 | 0 | 11 | 20 | 0.55 | 0.23 |
| 37 | 92 | 68 | 9 | 14 | 1 | 14 | 24 | 0.58 | 0.35 |
| 38 | 101 | 79 | 12 | 10 | 0 | 10 | 22 | 0.45 | 0.28 |
| 39 | 118 | 91 | 15 | 12 | 0 | 12 | 27 | 0.44 | 0.30 |
| 40 | 75 | 63 | 4 | 5 | 3 | 5 | 12 | 0.42 | 0.19 |
| 41 | 65 | 51 | 12 | 2 | 0 | 2 | 14 | 0.14 | 0.27 |
| 42 | 70 | 60 | 5 | 3 | 2 | 3 | 10 | 0.30 | 0.17 |
| 43 | 65 | 52 | 6 | 2 | 5 | 2 | 13 | 0.15 | 0.25 |
| 44 | 77 | 63 | 8 | 5 | 1 | 5 | 14 | 0.36 | 0.22 |
| 45 | 92 | 79 | 8 | 4 | 1 | 4 | 13 | 0.31 | 0.16 |
| 46 | 48 | 39 | 4 | 0 | 5 | 0 | 9 | 0.00 | 0.23 |
| 47 | 79 | 61 | 9 | 8 | 1 | 8 | 18 | 0.44 | 0.30 |
| 48 | 81 | 61 | 12 | 8 | 0 | 8 | 20 | 0.40 | 0.33 |
| 49 | 67 | 55 | 7 | 5 | 0 | 5 | 12 | 0.42 | 0.22 |
| 50 | 83 | 66 | 11 | 6 | 0 | 6 | 17 | 0.35 | 0.26 |
| 51 | 60 | 54 | 4 | 2 | 0 | 2 | 6 | 0.33 | 0.11 |
| 52 | 44 | 31 | 12 | 0 | 1 | 0 | 13 | 0.00 | 0.42 |
| 53 | 108 | 79 | 16 | 11 | 2 | 11 | 29 | 0.38 | 0.37 |
| 54 | 75 | 53 | 10 | 9 | 3 | 9 | 22 | 0.41 | 0.42 |
| 55 | 70 | 59 | 10 | 0 | 1 | 0 | 11 | 0.00 | 0.19 |
| 56 | 75 | 60 | 5 | 9 | 1 | 9 | 15 | 0.60 | 0.25 |
| 57 | 73 | 50 | 15 | 8 | 0 | 8 | 23 | 0.35 | 0.46 |
| 58 | 93 | 63 | 16 | 10 | 4 | 10 | 30 | 0.33 | 0.48 |
| 59 | 79 | 61 | 12 | 4 | 2 | 4 | 18 | 0.22 | 0.30 |
| 60 | 70 | 52 | 8 | 10 | 0 | 10 | 18 | 0.56 | 0.35 |
| 61 | 97 | 78 | 11 | 5 | 3 | 5 | 19 | 0.26 | 0.24 |
| 62 | 89 | 73 | 9 | 7 | 0 | 7 | 16 | 0.44 | 0.22 |
| 63 | 82 | 67 | 11 | 2 | 2 | 2 | 15 | 0.13 | 0.22 |
| 64 | 92 | 80 | 10 | 1 | 1 | 1 | 12 | 0.08 | 0.15 |
| 65 | 82 | 55 | 15 | 10 | 2 | 10 | 27 | 0.37 | 0.49 |
| 66 | 87 | 60 | 24 | 3 | 0 | 3 | 27 | 0.11 | 0.45 |
| 67 | 88 | 65 | 9 | 11 | 3 | 11 | 23 | 0.48 | 0.35 |
| 68 | | | | | | | | | |
| 69 | | | | | | | | | |
| 70 | | | | | | | | | |
| 71 | | | | | | | | | |
| 72 | | | | | | | | | |
| 73 | | | | | | | | | |
| 74 | | | | | | | | | |
| 75 | | | | | | | | | |
| 76 | | | | | | | | | |
| 77 | | | | | | | | | |
| 78 | | | | | | | | | |
| 79 | | | | | | | | | |
| 80 | | | | | | | | | |
| **TOTAL** | **5404** | **4218** | **675** | **426** | **85** | **426** | **1186** | 22.89 | 20.48 |
| **AVERAGE** | **80.7** | **63.0** | **10.1** | **6.4** | **1.3** | **6.4** | **17.7** | 0.34 | 0.31 |
| **SEM** | **1.9** | **1.6** | **0.4** | **0.4** | **0.2** | **0.4** | **0.6** | 0.02 | 0.02 |

#### p38 Inhibited

| # | TOTAL NUMBER OF CELLS | | ICM | | | | | Ratio of PrE/Total ICM | Ratio of Inner/ Outer cells |
|---|---|---|---|---|---|---|---|---|---|
| | Embryo | Double Negative | NANOG +ve | GATA4 +ve | NANOG/ GATA4 double +ve | PrE | Total | | |
| 1 | 15 | 15 | 0 | 0 | 0 | 0 | 0 | 0.00 | 0.00 |
| 2 | 28 | 27 | 1 | 0 | 0 | 0 | 1 | 0.00 | 0.04 |
| 3 | 45 | 40 | 4 | 1 | 0 | 1 | 5 | 0.20 | 0.13 |
| 4 | 65 | 53 | 8 | 3 | 1 | 3 | 12 | 0.25 | 0.23 |
| 5 | 41 | 37 | 4 | 0 | 0 | 0 | 4 | 0.00 | 0.11 |
| 6 | 38 | 35 | 3 | 0 | 0 | 0 | 3 | 0.00 | 0.09 |
| 7 | 35 | 25 | 9 | 1 | 0 | 1 | 10 | 0.10 | 0.40 |
| 8 | 49 | 41 | 7 | 1 | 0 | 1 | 8 | 0.13 | 0.20 |
| 9 | 29 | 26 | 3 | 0 | 0 | 0 | 3 | 0.00 | 0.12 |
| 10 | 27 | 22 | 5 | 0 | 0 | 0 | 5 | 0.00 | 0.23 |
| 11 | 34 | 29 | 5 | 0 | 0 | 0 | 5 | 0.00 | 0.17 |
| 12 | 32 | 30 | 2 | 0 | 0 | 0 | 2 | 0.00 | 0.07 |
| 13 | 69 | 49 | 16 | 4 | 0 | 4 | 20 | 0.20 | 0.41 |
| 14 | 61 | 51 | 6 | 4 | 0 | 4 | 10 | 0.40 | 0.20 |
| 15 | 78 | 67 | 5 | 6 | 0 | 6 | 11 | 0.55 | 0.16 |
| 16 | 41 | 40 | 1 | 0 | 0 | 0 | 1 | 0.00 | 0.03 |
| 17 | 37 | 31 | 4 | 0 | 2 | 0 | 6 | 0.00 | 0.19 |
| 18 | 54 | 45 | 5 | 3 | 1 | 3 | 9 | 0.33 | 0.20 |
| 19 | 41 | 37 | 3 | 0 | 1 | 0 | 4 | 0.00 | 0.11 |
| 20 | 37 | 34 | 2 | 0 | 1 | 0 | 3 | 0.00 | 0.09 |
| 21 | 31 | 19 | 11 | 1 | 0 | 1 | 12 | 0.08 | 0.63 |
| 22 | 31 | 29 | 2 | 0 | 0 | 0 | 2 | 0.00 | 0.07 |
| 23 | 27 | 17 | 10 | 0 | 0 | 0 | 10 | 0.00 | 0.59 |
| 24 | 22 | 18 | 4 | 0 | 0 | 0 | 4 | 0.00 | 0.22 |
| 25 | 36 | 30 | 6 | 0 | 0 | 0 | 6 | 0.00 | 0.20 |
| 26 | 59 | 50 | 8 | 1 | 0 | 1 | 9 | 0.11 | 0.18 |
| 27 | 53 | 46 | 5 | 1 | 1 | 1 | 7 | 0.14 | 0.15 |
| 28 | 49 | 36 | 9 | 4 | 0 | 4 | 13 | 0.31 | 0.36 |
| 29 | 57 | 51 | 4 | 2 | 0 | 2 | 6 | 0.33 | 0.12 |
| 30 | 70 | 56 | 8 | 6 | 0 | 6 | 14 | 0.43 | 0.25 |
| 31 | 62 | 50 | 12 | 0 | 0 | 0 | 12 | 0.00 | 0.24 |
| 32 | 43 | 36 | 5 | 2 | 0 | 2 | 7 | 0.29 | 0.19 |
| 33 | 33 | 27 | 5 | 1 | 0 | 1 | 6 | 0.17 | 0.22 |
| 34 | 22 | 19 | 3 | 0 | 0 | 0 | 3 | 0.00 | 0.16 |
| 35 | 39 | 33 | 5 | 1 | 0 | 1 | 6 | 0.17 | 0.18 |
| 36 | 67 | 60 | 3 | 2 | 2 | 2 | 7 | 0.29 | 0.12 |
| 37 | 60 | 40 | 15 | 5 | 0 | 5 | 20 | 0.25 | 0.50 |
| 38 | 45 | 41 | 4 | 0 | 0 | 0 | 4 | 0.00 | 0.10 |
| 39 | 30 | 26 | 3 | 1 | 0 | 1 | 4 | 0.25 | 0.15 |
| 40 | 65 | 56 | 6 | 1 | 2 | 1 | 9 | 0.11 | 0.16 |
| 41 | 68 | 60 | 8 | 0 | 0 | 0 | 8 | 0.00 | 0.13 |
| 42 | 45 | 34 | 11 | 0 | 0 | 0 | 11 | 0.00 | 0.32 |
| 43 | 58 | 54 | 4 | 0 | 0 | 0 | 4 | 0.00 | 0.07 |
| 44 | 57 | 49 | 5 | 3 | 0 | 3 | 8 | 0.38 | 0.16 |
| 45 | 70 | 65 | 3 | 0 | 2 | 0 | 5 | 0.00 | 0.08 |
| 46 | 37 | 35 | 2 | 0 | 0 | 0 | 2 | 0.00 | 0.06 |
| 47 | 75 | 67 | 6 | 2 | 0 | 2 | 8 | 0.25 | 0.12 |
| 48 | 29 | 18 | 11 | 0 | 0 | 0 | 11 | 0.00 | 0.61 |
| 49 | 50 | 43 | 6 | 1 | 0 | 1 | 7 | 0.14 | 0.16 |
| 50 | 25 | 16 | 9 | 0 | 0 | 0 | 9 | 0.00 | 0.56 |
| 51 | 32 | 29 | 3 | 0 | 0 | 0 | 3 | 0.00 | 0.10 |
| 52 | 25 | 22 | 3 | 0 | 0 | 0 | 3 | 0.00 | 0.14 |
| 53 | 29 | 25 | 4 | 0 | 0 | 0 | 4 | 0.00 | 0.16 |
| 54 | 54 | 48 | 6 | 0 | 0 | 0 | 6 | 0.00 | 0.13 |
| 55 | 37 | 30 | 7 | 0 | 0 | 0 | 7 | 0.00 | 0.23 |
| 56 | 28 | 27 | 1 | 0 | 0 | 0 | 1 | 0.00 | 0.04 |
| 57 | 12 | 12 | 0 | 0 | 0 | 0 | 0 | 0.00 | 0.00 |
| 58 | 31 | 27 | 2 | 2 | 0 | 2 | 4 | 0.50 | 0.15 |
| 59 | 50 | 38 | 11 | 0 | 1 | 0 | 12 | 0.00 | 0.32 |
| 60 | 66 | 58 | 8 | 0 | 0 | 0 | 8 | 0.00 | 0.14 |
| 61 | 44 | 38 | 6 | 0 | 0 | 0 | 6 | 0.00 | 0.16 |
| 62 | 50 | 46 | 4 | 0 | 0 | 0 | 4 | 0.00 | 0.09 |
| 63 | 39 | 35 | 4 | 0 | 0 | 0 | 4 | 0.00 | 0.11 |
| 64 | 76 | 72 | 4 | 0 | 0 | 0 | 4 | 0.00 | 0.06 |
| 65 | 34 | 31 | 3 | 0 | 0 | 0 | 3 | 0.00 | 0.10 |
| 66 | 55 | 52 | 3 | 0 | 0 | 0 | 3 | 0.00 | 0.06 |
| 67 | 39 | 34 | 5 | 0 | 0 | 0 | 5 | 0.00 | 0.15 |
| 68 | 31 | 24 | 7 | 0 | 0 | 0 | 7 | 0.00 | 0.29 |
| 69 | 38 | 21 | 17 | 0 | 0 | 0 | 17 | 0.00 | 0.81 |
| 70 | 38 | 29 | 9 | 0 | 0 | 0 | 9 | 0.00 | 0.31 |
| 71 | 90 | 84 | 5 | 1 | 0 | 1 | 6 | 0.17 | 0.07 |
| 72 | 45 | 37 | 8 | 0 | 0 | 0 | 8 | 0.00 | 0.22 |
| 73 | 27 | 16 | 11 | 0 | 0 | 0 | 11 | 0.00 | 0.69 |
| 74 | 40 | 35 | 5 | 0 | 0 | 0 | 5 | 0.00 | 0.14 |
| 75 | 37 | 30 | 5 | 0 | 2 | 0 | 7 | 0.00 | 0.23 |
| 76 | 52 | 43 | 9 | 0 | 0 | 0 | 9 | 0.00 | 0.21 |
| 77 | 45 | 34 | 11 | 0 | 0 | 0 | 11 | 0.00 | 0.32 |
| 78 | | | | | | | | | |
| 79 | | | | | | | | | |
| 80 | | | | | | | | | |
| **TOTAL** | **3415** | **2892** | **447** | **60** | **16** | **60** | **523** | 6.51 | 15.63 |
| **AVERAGE** | **44.4** | **37.6** | **5.8** | **0.8** | **0.2** | **0.8** | **6.8** | 0.08 | 0.20 |
| **SEM** | **1.8** | **1.7** | **0.4** | **0.2** | **0.1** | **0.2** | **0.5** | 0.02 | 0.02 |

Supplementary table S1 (contd.)

### KSOM with amino acids

#### DMSO

| # | TOTAL NUMBER OF CELLS: Embryo | Double Negative | ICM: NANOG +ve | GATA4 +ve | NANOG/ GATA4 double +ve | PrE | Total | Ratio of PrE/Total ICM | Ratio of Inner/ Outer cells |
|---|---|---|---|---|---|---|---|---|---|
| 1 | 52 | 34 | 17 | 0 | 1 | 0 | 18 | 0.00 | 0.53 |
| 2 | 84 | 67 | 9 | 8 | 0 | 8 | 17 | 0.47 | 0.25 |
| 3 | 68 | 51 | 15 | 2 | 0 | 2 | 17 | 0.12 | 0.33 |
| 4 | 83 | 69 | 7 | 7 | 0 | 7 | 14 | 0.50 | 0.20 |
| 5 | 66 | 57 | 8 | 1 | 0 | 1 | 9 | 0.11 | 0.16 |
| 6 | 87 | 73 | 9 | 5 | 0 | 5 | 14 | 0.36 | 0.19 |
| 7 | 59 | 50 | 8 | 1 | 0 | 1 | 9 | 0.11 | 0.18 |
| 8 | 74 | 62 | 11 | 1 | 0 | 1 | 12 | 0.08 | 0.19 |
| 9 | 81 | 59 | 16 | 4 | 2 | 4 | 22 | 0.18 | 0.37 |
| 10 | 66 | 48 | 10 | 7 | 1 | 7 | 18 | 0.39 | 0.38 |
| 11 | 75 | 61 | 7 | 7 | 0 | 7 | 14 | 0.50 | 0.23 |
| 12 | 61 | 48 | 7 | 6 | 0 | 6 | 13 | 0.46 | 0.27 |
| 13 | 78 | 64 | 10 | 3 | 1 | 3 | 14 | 0.21 | 0.22 |
| 14 | 74 | 60 | 6 | 8 | 0 | 8 | 14 | 0.57 | 0.23 |
| 15 | 71 | 48 | 11 | 12 | 0 | 12 | 23 | 0.52 | 0.48 |
| 16 | 78 | 62 | 7 | 8 | 1 | 8 | 16 | 0.50 | 0.26 |
| 17 | 79 | 66 | 6 | 4 | 3 | 4 | 13 | 0.31 | 0.20 |
| 18 | 84 | 61 | 10 | 12 | 1 | 12 | 23 | 0.52 | 0.38 |
| 19 | 44 | 35 | 4 | 3 | 2 | 3 | 9 | 0.33 | 0.26 |
| 20 | 91 | 70 | 14 | 7 | 0 | 7 | 21 | 0.33 | 0.30 |
| 21 | 76 | 67 | 5 | 3 | 1 | 3 | 9 | 0.33 | 0.13 |
| 22 | 78 | 63 | 10 | 5 | 0 | 5 | 15 | 0.33 | 0.24 |
| 23 | 93 | 75 | 10 | 7 | 1 | 7 | 18 | 0.39 | 0.24 |
| 24 | 96 | 66 | 16 | 14 | 0 | 14 | 30 | 0.47 | 0.45 |
| 25 | 76 | 48 | 11 | 17 | 0 | 17 | 28 | 0.61 | 0.58 |
| 26 | 80 | 59 | 8 | 13 | 0 | 13 | 21 | 0.62 | 0.36 |
| 27 | 90 | 80 | 5 | 5 | 0 | 5 | 10 | 0.50 | 0.13 |
| 28 | 72 | 57 | 10 | 5 | 0 | 5 | 15 | 0.33 | 0.26 |
| 29 | 83 | 64 | 12 | 7 | 0 | 7 | 19 | 0.37 | 0.30 |
| 30 | 95 | 69 | 13 | 12 | 1 | 12 | 26 | 0.46 | 0.38 |
| 31 | 82 | 62 | 10 | 9 | 1 | 9 | 20 | 0.45 | 0.32 |
| 32 | 69 | 58 | 8 | 2 | 1 | 2 | 11 | 0.18 | 0.19 |
| 33 | 88 | 71 | 13 | 4 | 0 | 4 | 17 | 0.24 | 0.24 |
| 34 | 77 | 66 | 3 | 8 | 0 | 8 | 11 | 0.73 | 0.17 |
| 35 | 84 | 60 | 9 | 15 | 0 | 15 | 24 | 0.63 | 0.40 |
| 36 | 86 | 62 | 14 | 9 | 1 | 9 | 24 | 0.38 | 0.39 |
| 37 | 89 | 70 | 13 | 6 | 0 | 6 | 19 | 0.32 | 0.27 |
| 38 | 72 | 58 | 11 | 1 | 2 | 1 | 14 | 0.07 | 0.24 |
| 39 | 58 | 48 | 6 | 4 | 0 | 4 | 10 | 0.40 | 0.21 |
| 40 | 73 | 52 | 6 | 15 | 0 | 15 | 21 | 0.71 | 0.40 |
| 41 | 85 | 63 | 15 | 5 | 2 | 5 | 22 | 0.23 | 0.35 |
| 42 | 90 | 67 | 7 | 15 | 1 | 15 | 23 | 0.65 | 0.34 |
| 43 | 85 | 67 | 9 | 7 | 2 | 7 | 18 | 0.39 | 0.27 |
| 44 | 117 | 98 | 11 | 7 | 1 | 7 | 19 | 0.37 | 0.19 |
| 45 | 84 | 74 | 6 | 4 | 0 | 4 | 10 | 0.40 | 0.14 |
| 46 | 88 | 69 | 10 | 8 | 1 | 8 | 19 | 0.42 | 0.28 |
| 47 | 107 | 89 | 11 | 7 | 0 | 7 | 18 | 0.39 | 0.20 |
| 48 | 98 | 68 | 12 | 16 | 2 | 16 | 30 | 0.53 | 0.44 |
| 49 | 94 | 81 | 11 | 1 | 1 | 1 | 13 | 0.08 | 0.16 |
| 50 | 90 | 67 | 16 | 7 | 0 | 7 | 23 | 0.30 | 0.34 |
| 51 | 80 | 60 | 10 | 8 | 2 | 8 | 20 | 0.40 | 0.33 |
| 52 | 79 | 57 | 8 | 12 | 2 | 12 | 22 | 0.55 | 0.39 |
| 53 | 93 | 71 | 19 | 3 | 0 | 3 | 22 | 0.14 | 0.31 |
| 54 | 66 | 45 | 18 | 3 | 0 | 3 | 21 | 0.14 | 0.47 |
| 55 | 101 | 74 | 14 | 11 | 2 | 11 | 27 | 0.41 | 0.36 |
| 56 | 72 | 55 | 12 | 5 | 0 | 5 | 17 | 0.29 | 0.31 |
| 57 | 83 | 57 | 15 | 9 | 2 | 9 | 26 | 0.35 | 0.46 |
| 58 | 71 | 40 | 21 | 5 | 5 | 5 | 31 | 0.16 | 0.78 |
| 59 | 92 | 73 | 8 | 7 | 4 | 7 | 19 | 0.37 | 0.26 |
| 60 | 78 | 58 | 11 | 3 | 6 | 3 | 20 | 0.15 | 0.34 |
| 61 | 80 | 60 | 13 | 7 | 0 | 7 | 20 | 0.35 | 0.33 |
| 62 | 77 | 56 | 10 | 11 | 0 | 11 | 21 | 0.52 | 0.38 |
| 63 | 60 | 40 | 20 | 0 | 0 | 0 | 20 | 0.00 | 0.50 |
| 64 | 78 | 69 | 6 | 3 | 0 | 3 | 9 | 0.33 | 0.13 |
| 65 | 95 | 75 | 8 | 9 | 3 | 9 | 20 | 0.45 | 0.27 |
| 66 | 106 | 88 | 13 | 4 | 1 | 4 | 18 | 0.22 | 0.20 |
| 67 | 80 | 62 | 8 | 8 | 2 | 8 | 18 | 0.44 | 0.29 |
| 68 | 111 | 82 | 12 | 17 | 0 | 17 | 29 | 0.59 | 0.35 |
| 69 | 110 | 88 | 10 | 12 | 0 | 12 | 22 | 0.55 | 0.25 |
| 70 | 76 | 59 | 10 | 7 | 0 | 7 | 17 | 0.41 | 0.29 |
| 71 | 96 | 76 | 10 | 10 | 0 | 10 | 20 | 0.50 | 0.26 |
| 72 | | | | | | | | | |
| 73 | | | | | | | | | |
| 74 | | | | | | | | | |
| 75 | | | | | | | | | |
| 76 | | | | | | | | | |
| 77 | | | | | | | | | |
| 78 | | | | | | | | | |
| 79 | | | | | | | | | |
| 80 | | | | | | | | | |
| **TOTAL** | **5794** | **4488** | **749** | **498** | **59** | **498** | **1306** | 26.17 | 21.48 |
| **AVERAGE** | **81.6** | **63.2** | **10.5** | **7.0** | **0.8** | **7.0** | **18.4** | 0.37 | 0.30 |
| **SEM** | **1.6** | **1.5** | **0.4** | **0.5** | **0.1** | **0.5** | **0.7** | 0.02 | 0.01 |

#### p38 Inhibited

| # | TOTAL NUMBER OF CELLS: Embryo | Double Negative | ICM: NANOG +ve | GATA4 +ve | NANOG/ GATA4 double +ve | PrE | Total | Ratio of PrE/Total ICM | Ratio of Inner/ Outer cells |
|---|---|---|---|---|---|---|---|---|---|
| 1 | 79 | 71 | 7 | 0 | 1 | 0 | 8 | 0.00 | 0.11 |
| 2 | 66 | 55 | 8 | 3 | 0 | 3 | 11 | 0.27 | 0.20 |
| 3 | 63 | 49 | 11 | 3 | 0 | 3 | 14 | 0.21 | 0.29 |
| 4 | 59 | 53 | 5 | 0 | 1 | 0 | 6 | 0.00 | 0.11 |
| 5 | 52 | 45 | 6 | 1 | 0 | 1 | 7 | 0.14 | 0.16 |
| 6 | 54 | 47 | 5 | 0 | 2 | 0 | 7 | 0.00 | 0.15 |
| 7 | 56 | 49 | 6 | 1 | 0 | 1 | 7 | 0.14 | 0.14 |
| 8 | 46 | 38 | 8 | 0 | 0 | 0 | 8 | 0.00 | 0.21 |
| 9 | 47 | 42 | 5 | 0 | 0 | 0 | 5 | 0.00 | 0.12 |
| 10 | 53 | 43 | 6 | 3 | 1 | 3 | 10 | 0.30 | 0.23 |
| 11 | 36 | 27 | 6 | 1 | 2 | 1 | 9 | 0.11 | 0.33 |
| 12 | 70 | 56 | 10 | 3 | 1 | 3 | 14 | 0.21 | 0.25 |
| 13 | 55 | 46 | 7 | 2 | 0 | 2 | 9 | 0.22 | 0.20 |
| 14 | 46 | 38 | 5 | 0 | 3 | 0 | 8 | 0.00 | 0.21 |
| 15 | 56 | 49 | 5 | 1 | 1 | 1 | 7 | 0.14 | 0.14 |
| 16 | 47 | 37 | 9 | 1 | 0 | 1 | 10 | 0.10 | 0.27 |
| 17 | 62 | 52 | 7 | 3 | 0 | 3 | 10 | 0.30 | 0.19 |
| 18 | 57 | 42 | 14 | 1 | 0 | 1 | 15 | 0.07 | 0.36 |
| 19 | 55 | 44 | 9 | 1 | 1 | 1 | 11 | 0.09 | 0.25 |
| 20 | 51 | 42 | 9 | 0 | 0 | 0 | 9 | 0.00 | 0.21 |
| 21 | 53 | 42 | 8 | 2 | 1 | 2 | 11 | 0.18 | 0.26 |
| 22 | 78 | 60 | 9 | 8 | 1 | 8 | 18 | 0.44 | 0.30 |
| 23 | 61 | 56 | 5 | 0 | 0 | 0 | 5 | 0.00 | 0.09 |
| 24 | 45 | 40 | 5 | 0 | 0 | 0 | 5 | 0.00 | 0.13 |
| 25 | 51 | 37 | 14 | 0 | 0 | 0 | 14 | 0.00 | 0.38 |
| 26 | 45 | 34 | 10 | 1 | 0 | 1 | 11 | 0.09 | 0.32 |
| 27 | 53 | 46 | 5 | 1 | 1 | 1 | 7 | 0.14 | 0.15 |
| 28 | 62 | 51 | 8 | 3 | 0 | 3 | 11 | 0.27 | 0.22 |
| 29 | 55 | 40 | 9 | 5 | 1 | 5 | 15 | 0.33 | 0.38 |
| 30 | 58 | 50 | 6 | 2 | 0 | 2 | 8 | 0.25 | 0.16 |
| 31 | 64 | 54 | 7 | 0 | 3 | 0 | 10 | 0.00 | 0.19 |
| 32 | 78 | 65 | 11 | 2 | 0 | 2 | 13 | 0.15 | 0.20 |
| 33 | 65 | 48 | 9 | 8 | 0 | 8 | 17 | 0.47 | 0.35 |
| 34 | 35 | 29 | 6 | 0 | 0 | 0 | 6 | 0.00 | 0.21 |
| 35 | 73 | 53 | 15 | 5 | 0 | 5 | 20 | 0.25 | 0.38 |
| 36 | 71 | 48 | 23 | 0 | 0 | 0 | 23 | 0.00 | 0.48 |
| 37 | 77 | 57 | 10 | 10 | 0 | 10 | 20 | 0.50 | 0.35 |
| 38 | 61 | 50 | 10 | 1 | 0 | 1 | 11 | 0.09 | 0.22 |
| 39 | 75 | 65 | 7 | 3 | 0 | 3 | 10 | 0.30 | 0.15 |
| 40 | 62 | 43 | 16 | 3 | 0 | 3 | 19 | 0.16 | 0.44 |
| 41 | 61 | 47 | 12 | 2 | 0 | 2 | 14 | 0.14 | 0.30 |
| 42 | 79 | 66 | 9 | 4 | 0 | 4 | 13 | 0.31 | 0.20 |
| 43 | 23 | 18 | 5 | 0 | 0 | 0 | 5 | 0.00 | 0.28 |
| 44 | 68 | 54 | 9 | 4 | 1 | 4 | 14 | 0.29 | 0.26 |
| 45 | 68 | 59 | 5 | 4 | 0 | 4 | 9 | 0.44 | 0.15 |
| 46 | 57 | 43 | 9 | 5 | 0 | 5 | 14 | 0.36 | 0.33 |
| 47 | 63 | 50 | 13 | 0 | 0 | 0 | 13 | 0.00 | 0.26 |
| 48 | 64 | 59 | 5 | 0 | 0 | 0 | 5 | 0.00 | 0.08 |
| 49 | 66 | 55 | 10 | 1 | 0 | 1 | 11 | 0.09 | 0.20 |
| 50 | 52 | 44 | 4 | 3 | 1 | 3 | 8 | 0.38 | 0.18 |
| 51 | 68 | 61 | 7 | 0 | 0 | 0 | 7 | 0.00 | 0.11 |
| 52 | 66 | 50 | 15 | 1 | 0 | 1 | 16 | 0.06 | 0.32 |
| 53 | 60 | 49 | 7 | 2 | 2 | 2 | 11 | 0.18 | 0.22 |
| 54 | 49 | 37 | 12 | 0 | 0 | 0 | 12 | 0.00 | 0.32 |
| 55 | 57 | 50 | 7 | 0 | 0 | 0 | 7 | 0.00 | 0.14 |
| 56 | 85 | 60 | 16 | 8 | 1 | 8 | 25 | 0.32 | 0.42 |
| 57 | 63 | 54 | 8 | 0 | 1 | 0 | 9 | 0.00 | 0.17 |
| 58 | 71 | 60 | 11 | 0 | 0 | 0 | 11 | 0.00 | 0.18 |
| 59 | 53 | 34 | 18 | 0 | 1 | 0 | 19 | 0.00 | 0.56 |
| 60 | 51 | 48 | 3 | 0 | 0 | 0 | 3 | 0.00 | 0.06 |
| 61 | 49 | 39 | 9 | 1 | 0 | 1 | 10 | 0.10 | 0.26 |
| 62 | 35 | 27 | 7 | 1 | 0 | 1 | 8 | 0.13 | 0.30 |
| 63 | 45 | 28 | 17 | 0 | 0 | 0 | 17 | 0.00 | 0.61 |
| 64 | 71 | 59 | 8 | 1 | 3 | 1 | 12 | 0.08 | 0.20 |
| 65 | 55 | 49 | 5 | 1 | 0 | 1 | 6 | 0.17 | 0.12 |
| 66 | 59 | 29 | 30 | 0 | 0 | 0 | 30 | 0.00 | 1.03 |
| 67 | 43 | 24 | 19 | 0 | 0 | 0 | 19 | 0.00 | 0.79 |
| 68 | 52 | 44 | 5 | 2 | 1 | 2 | 8 | 0.25 | 0.18 |
| 69 | 55 | 34 | 20 | 0 | 1 | 0 | 21 | 0.00 | 0.62 |
| 70 | 66 | 56 | 10 | 0 | 0 | 0 | 10 | 0.00 | 0.18 |
| 71 | 65 | 50 | 11 | 4 | 0 | 4 | 15 | 0.27 | 0.30 |
| 72 | 62 | 49 | 11 | 2 | 0 | 2 | 13 | 0.15 | 0.27 |
| 73 | 48 | 36 | 12 | 0 | 0 | 0 | 12 | 0.00 | 0.33 |
| 74 | 85 | 69 | 6 | 10 | 0 | 10 | 16 | 0.63 | 0.23 |
| 75 | 73 | 56 | 14 | 1 | 2 | 1 | 17 | 0.06 | 0.30 |
| 76 | 76 | 54 | 11 | 11 | 0 | 11 | 22 | 0.50 | 0.41 |
| 77 | | | | | | | | | |
| 78 | | | | | | | | | |
| 79 | | | | | | | | | |
| 80 | | | | | | | | | |
| **TOTAL** | **4495** | **3594** | **721** | **146** | **34** | **146** | **901** | 10.86 | 20.37 |
| **AVERAGE** | **59.1** | **47.3** | **9.5** | **1.9** | **0.4** | **1.9** | **11.9** | 0.14 | 0.27 |
| **SEM** | **1.4** | **1.2** | **0.5** | **0.3** | **0.1** | **0.3** | **0.6** | 0.02 | 0.02 |

Supplementary table S2

### KSOM + 1mM NAC

#### DMSO

| # | TOTAL NUMBER OF CELLS | | | | | | | Ratio of PrE/Total ICM | Ratio of Inner/ Outer cells |
|---|---|---|---|---|---|---|---|---|---|
| | Embryo | Double Negative | ICM | | | | | | |
| | | | NANOG +ve | GATA4 +ve | NANOG/ GATA4 double +ve | PrE | Total | | |
| 1 | 82 | 63 | 8 | 11 | 0 | 11 | 19 | 0.58 | 0.30 |
| 2 | 65 | 63 | 2 | 0 | 0 | 0 | 2 | 0.00 | 0.03 |
| 3 | 38 | 28 | 9 | 0 | 1 | 0 | 10 | 0.00 | 0.36 |
| 4 | 70 | 49 | 11 | 10 | 0 | 10 | 21 | 0.48 | 0.43 |
| 5 | 70 | 60 | 6 | 4 | 0 | 4 | 10 | 0.40 | 0.17 |
| 6 | 71 | 53 | 10 | 8 | 0 | 8 | 18 | 0.44 | 0.34 |
| 7 | 89 | 73 | 10 | 6 | 0 | 6 | 16 | 0.38 | 0.22 |
| 8 | 83 | 70 | 5 | 6 | 2 | 6 | 13 | 0.46 | 0.19 |
| 9 | 95 | 80 | 7 | 8 | 0 | 8 | 15 | 0.53 | 0.19 |
| 10 | 96 | 82 | 9 | 5 | 0 | 5 | 14 | 0.36 | 0.17 |
| 11 | 88 | 73 | 9 | 6 | 0 | 6 | 15 | 0.40 | 0.21 |
| 12 | 74 | 68 | 3 | 3 | 0 | 3 | 6 | 0.50 | 0.09 |
| 13 | 85 | 60 | 12 | 13 | 0 | 13 | 25 | 0.52 | 0.42 |
| 14 | 98 | 69 | 16 | 13 | 0 | 13 | 29 | 0.45 | 0.42 |
| 15 | 111 | 79 | 21 | 11 | 0 | 11 | 32 | 0.34 | 0.41 |
| 16 | 80 | 65 | 12 | 3 | 0 | 3 | 15 | 0.20 | 0.23 |
| 17 | 77 | 57 | 14 | 5 | 1 | 5 | 20 | 0.25 | 0.35 |
| 18 | 97 | 75 | 13 | 8 | 1 | 8 | 22 | 0.36 | 0.29 |
| 19 | 73 | 51 | 17 | 2 | 3 | 2 | 22 | 0.09 | 0.43 |
| 20 | 70 | 43 | 14 | 10 | 3 | 10 | 27 | 0.37 | 0.63 |
| 21 | 91 | 68 | 13 | 10 | 0 | 10 | 23 | 0.43 | 0.34 |
| 22 | 62 | 50 | 7 | 2 | 3 | 2 | 12 | 0.17 | 0.24 |
| 23 | 75 | 47 | 18 | 9 | 1 | 9 | 28 | 0.32 | 0.60 |
| 24 | 91 | 73 | 10 | 7 | 1 | 7 | 18 | 0.39 | 0.25 |
| 25 | 64 | 52 | 6 | 6 | 0 | 6 | 12 | 0.50 | 0.23 |
| 26 | 84 | 71 | 8 | 4 | 1 | 4 | 13 | 0.31 | 0.18 |
| 27 | 77 | 58 | 11 | 8 | 0 | 8 | 19 | 0.42 | 0.33 |
| 28 | 84 | 70 | 9 | 5 | 0 | 5 | 14 | 0.36 | 0.20 |
| 29 | | | | | | | | | |
| 30 | | | | | | | | | |
| 31 | | | | | | | | | |
| 32 | | | | | | | | | |
| 33 | | | | | | | | | |
| 34 | | | | | | | | | |
| 35 | | | | | | | | | |
| 36 | | | | | | | | | |
| 37 | | | | | | | | | |
| 38 | | | | | | | | | |
| 39 | | | | | | | | | |
| 40 | | | | | | | | | |
| **TOTAL** | **2240** | **1750** | **290** | **183** | **17** | **183** | **490** | 10.01 | 8.22 |
| **AVERAGE** | **80.0** | **62.5** | **10.4** | **6.5** | **0.6** | **6.5** | **17.5** | 0.36 | 0.29 |
| **SEM** | **2.7** | **2.4** | **0.8** | **0.7** | **0.2** | **0.7** | **1.3** | 0.03 | 0.03 |

#### p38 Inhibited

| # | TOTAL NUMBER OF CELLS | | | | | | | Ratio of PrE/Total ICM | Ratio of Inner/ Outer cells |
|---|---|---|---|---|---|---|---|---|---|
| | Embryo | Double Negative | ICM | | | | | | |
| | | | NANOG +ve | GATA4 +ve | NANOG/ GATA4 double +ve | PrE | Total | | |
| 1 | 43 | 41 | 2 | 0 | 0 | 0 | 2 | 0.00 | 0.05 |
| 2 | 42 | 36 | 4 | 2 | 0 | 2 | 6 | 0.33 | 0.17 |
| 3 | 46 | 43 | 3 | 0 | 0 | 0 | 3 | 0.00 | 0.07 |
| 4 | 63 | 56 | 5 | 2 | 0 | 2 | 7 | 0.29 | 0.13 |
| 5 | 58 | 48 | 10 | 0 | 0 | 0 | 10 | 0.00 | 0.21 |
| 6 | 27 | 16 | 11 | 0 | 0 | 0 | 11 | 0.00 | 0.69 |
| 7 | 38 | 36 | 2 | 0 | 0 | 0 | 2 | 0.00 | 0.06 |
| 8 | 56 | 48 | 8 | 0 | 0 | 0 | 8 | 0.00 | 0.17 |
| 9 | 34 | 25 | 6 | 3 | 0 | 3 | 9 | 0.33 | 0.36 |
| 10 | 51 | 37 | 9 | 5 | 0 | 5 | 14 | 0.36 | 0.38 |
| 11 | 29 | 11 | 18 | 0 | 0 | 0 | 18 | 0.00 | 1.64 |
| 12 | 48 | 43 | 3 | 2 | 0 | 2 | 5 | 0.40 | 0.12 |
| 13 | 34 | 30 | 4 | 0 | 0 | 0 | 4 | 0.00 | 0.13 |
| 14 | 35 | 29 | 6 | 0 | 0 | 0 | 6 | 0.00 | 0.21 |
| 15 | 52 | 46 | 2 | 3 | 1 | 3 | 6 | 0.50 | 0.13 |
| 16 | 58 | 47 | 9 | 2 | 0 | 2 | 11 | 0.18 | 0.23 |
| 17 | 41 | 34 | 5 | 2 | 0 | 2 | 7 | 0.29 | 0.21 |
| 18 | 67 | 53 | 8 | 6 | 0 | 6 | 14 | 0.43 | 0.26 |
| 19 | 67 | 62 | 5 | 0 | 0 | 0 | 5 | 0.00 | 0.08 |
| 20 | 70 | 54 | 14 | 2 | 0 | 2 | 16 | 0.13 | 0.30 |
| 21 | 77 | 63 | 9 | 5 | 0 | 5 | 14 | 0.36 | 0.22 |
| 22 | 43 | 29 | 11 | 1 | 2 | 1 | 14 | 0.07 | 0.48 |
| 23 | 58 | 50 | 6 | 0 | 2 | 0 | 8 | 0.00 | 0.16 |
| 24 | 92 | 78 | 9 | 4 | 1 | 4 | 14 | 0.29 | 0.18 |
| 25 | 71 | 61 | 5 | 5 | 0 | 5 | 10 | 0.50 | 0.16 |
| 26 | 55 | 46 | 7 | 2 | 0 | 2 | 9 | 0.22 | 0.20 |
| 27 | 47 | 40 | 6 | 0 | 1 | 0 | 7 | 0.00 | 0.18 |
| 28 | 78 | 67 | 11 | 0 | 0 | 0 | 11 | 0.00 | 0.16 |
| 29 | 83 | 81 | 2 | 0 | 0 | 0 | 2 | 0.00 | 0.02 |
| 30 | 81 | 70 | 7 | 4 | 0 | 4 | 11 | 0.36 | 0.16 |
| 31 | 47 | 46 | 1 | 0 | 0 | 0 | 1 | 0.00 | 0.02 |
| 32 | 45 | 38 | 7 | 0 | 0 | 0 | 7 | 0.00 | 0.18 |
| 33 | 75 | 67 | 4 | 4 | 0 | 4 | 8 | 0.50 | 0.12 |
| 34 | 65 | 61 | 2 | 2 | 0 | 2 | 4 | 0.50 | 0.07 |
| 35 | 60 | 51 | 9 | 0 | 0 | 0 | 9 | 0.00 | 0.18 |
| 36 | 70 | 63 | 4 | 3 | 0 | 3 | 7 | 0.43 | 0.11 |
| 37 | 60 | 51 | 9 | 0 | 0 | 0 | 9 | 0.00 | 0.18 |
| 38 | 59 | 59 | 0 | 0 | 0 | 0 | 0 | 0.00 | 0.00 |
| 39 | | | | | | | | | |
| 40 | | | | | | | | | |
| **TOTAL** | **2125** | **1816** | **243** | **59** | **7** | **59** | **309** | 6.46 | 8.35 |
| **AVERAGE** | **55.9** | **47.8** | **6.4** | **1.6** | **0.2** | **1.6** | **8.1** | 0.17 | 0.22 |
| **SEM** | **2.6** | **2.6** | **0.6** | **0.3** | **0.1** | **0.3** | **0.7** | 0.03 | 0.04 |

### KSOM + 10mM NAC

#### DMSO

| # | TOTAL NUMBER OF CELLS | | | | | | | Ratio of PrE/Total ICM | Ratio of Inner/ Outer cells |
|---|---|---|---|---|---|---|---|---|---|
| | Embryo | Double Negative | ICM | | | | | | |
| | | | NANOG +ve | GATA4 +ve | NANOG/ GATA4 double +ve | PrE | Total | | |
| 1 | 57 | 39 | 10 | 8 | 0 | 8 | 18 | 0.44 | 0.46 |
| 2 | 75 | 53 | 16 | 6 | 0 | 6 | 22 | 0.27 | 0.42 |
| 3 | 74 | 58 | 9 | 7 | 0 | 7 | 16 | 0.44 | 0.28 |
| 4 | 77 | 59 | 15 | 3 | 0 | 3 | 18 | 0.17 | 0.31 |
| 5 | 43 | 38 | 5 | 0 | 0 | 0 | 5 | 0.00 | 0.13 |
| 6 | 57 | 48 | 3 | 5 | 1 | 5 | 9 | 0.56 | 0.19 |
| 7 | 51 | 40 | 11 | 0 | 0 | 0 | 11 | 0.00 | 0.28 |
| 8 | 55 | 41 | 10 | 4 | 0 | 4 | 14 | 0.29 | 0.34 |
| 9 | 103 | 83 | 10 | 8 | 2 | 8 | 20 | 0.40 | 0.24 |
| 10 | 81 | 60 | 10 | 11 | 0 | 11 | 21 | 0.52 | 0.35 |
| 11 | 82 | 61 | 9 | 11 | 1 | 11 | 21 | 0.52 | 0.34 |
| 12 | 95 | 78 | 7 | 10 | 0 | 10 | 17 | 0.59 | 0.22 |
| 13 | 83 | 72 | 8 | 3 | 0 | 3 | 11 | 0.27 | 0.15 |
| 14 | 93 | 75 | 9 | 9 | 0 | 9 | 18 | 0.50 | 0.24 |
| 15 | 83 | 66 | 9 | 8 | 0 | 8 | 17 | 0.47 | 0.26 |
| 16 | 102 | 86 | 7 | 9 | 0 | 9 | 16 | 0.56 | 0.19 |
| 17 | 72 | 48 | 12 | 12 | 0 | 12 | 24 | 0.50 | 0.50 |
| 18 | | | | | | | | | |
| 19 | | | | | | | | | |
| 20 | | | | | | | | | |
| **TOTAL** | **1283** | **1005** | **160** | **114** | **4** | **114** | **278** | 6.50 | 4.88 |
| **AVERAGE** | **75.5** | **59.1** | **9.4** | **6.7** | **0.2** | **6.7** | **16.4** | 0.38 | 0.29 |
| **SEM** | **4.3** | **3.8** | **0.8** | **0.9** | **0.1** | **0.9** | **1.2** | 0.05 | 0.03 |

#### p38 Inhibited

| # | TOTAL NUMBER OF CELLS | | | | | | | Ratio of PrE/Total ICM | Ratio of Inner/ Outer cells |
|---|---|---|---|---|---|---|---|---|---|
| | Embryo | Double Negative | ICM | | | | | | |
| | | | NANOG +ve | GATA4 +ve | NANOG/ GATA4 double +ve | PrE | Total | | |
| 1 | 30 | 27 | 3 | 0 | 0 | 0 | 3 | 0.00 | 0.11 |
| 2 | 61 | 56 | 2 | 3 | 0 | 3 | 5 | 0.60 | 0.09 |
| 3 | 45 | 35 | 6 | 4 | 0 | 4 | 10 | 0.40 | 0.29 |
| 4 | 52 | 37 | 9 | 2 | 4 | 2 | 15 | 0.13 | 0.41 |
| 5 | 40 | 32 | 4 | 4 | 0 | 4 | 8 | 0.50 | 0.25 |
| 6 | 50 | 44 | 6 | 0 | 0 | 0 | 6 | 0.00 | 0.14 |
| 7 | 49 | 45 | 4 | 0 | 0 | 0 | 4 | 0.00 | 0.09 |
| 8 | 52 | 39 | 6 | 6 | 1 | 6 | 13 | 0.46 | 0.33 |
| 9 | 68 | 55 | 9 | 4 | 0 | 4 | 13 | 0.31 | 0.24 |
| 10 | 90 | 74 | 8 | 8 | 0 | 8 | 16 | 0.50 | 0.22 |
| 11 | 69 | 61 | 8 | 0 | 0 | 0 | 8 | 0.00 | 0.13 |
| 12 | 78 | 59 | 13 | 6 | 0 | 6 | 19 | 0.32 | 0.32 |
| 13 | 62 | 52 | 10 | 0 | 0 | 0 | 10 | 0.00 | 0.19 |
| 14 | 47 | 39 | 6 | 2 | 0 | 2 | 8 | 0.25 | 0.21 |
| 15 | 81 | 67 | 8 | 6 | 0 | 6 | 14 | 0.43 | 0.21 |
| 16 | 72 | 64 | 7 | 1 | 0 | 1 | 8 | 0.13 | 0.13 |
| 17 | 52 | 45 | 5 | 2 | 0 | 2 | 7 | 0.29 | 0.16 |
| 18 | 36 | 25 | 8 | 3 | 0 | 3 | 11 | 0.27 | 0.44 |
| 19 | | | | | | | | | |
| 20 | | | | | | | | | |
| **TOTAL** | **1034** | **856** | **122** | **51** | **5** | **51** | **178** | 4.58 | 3.93 |
| **AVERAGE** | **57.4** | **47.6** | **6.8** | **2.8** | **0.3** | **2.8** | **9.9** | 0.25 | 0.22 |
| **SEM** | **3.8** | **3.3** | **0.6** | **0.6** | **0.2** | **0.6** | **1.0** | 0.05 | 0.02 |

Supplementary table S2 (contd.)

### KSOM+AA + 1mM NAC

#### DMSO

| # | TOTAL NUMBER OF CELLS: Embryo | Double Negative | ICM: NANOG +ve | ICM: GATA4 +ve | ICM: NANOG/GATA4 double +ve | ICM: PrE | ICM: Total | Ratio of PrE/Total ICM | Ratio of Inner/Outer cells |
|---|---|---|---|---|---|---|---|---|---|
| 1 | 97 | 75 | 12 | 9 | 1 | 9 | 22 | 0.41 | 0.29 |
| 2 | 87 | 77 | 7 | 1 | 2 | 1 | 10 | 0.10 | 0.13 |
| 3 | 49 | 33 | 10 | 3 | 3 | 3 | 16 | 0.19 | 0.48 |
| 4 | 45 | 30 | 11 | 1 | 3 | 1 | 15 | 0.07 | 0.50 |
| 5 | 68 | 56 | 9 | 2 | 1 | 2 | 12 | 0.17 | 0.21 |
| 6 | 78 | 50 | 17 | 11 | 0 | 11 | 28 | 0.39 | 0.56 |
| 7 | 78 | 62 | 10 | 5 | 1 | 5 | 16 | 0.31 | 0.26 |
| 8 | 56 | 53 | 3 | 0 | 0 | 0 | 3 | 0.00 | 0.06 |
| 9 | 68 | 45 | 16 | 5 | 2 | 5 | 23 | 0.22 | 0.51 |
| 10 | 91 | 73 | 6 | 12 | 0 | 12 | 18 | 0.67 | 0.25 |
| 11 | 78 | 57 | 13 | 8 | 0 | 8 | 21 | 0.38 | 0.37 |
| 12 | 89 | 68 | 5 | 12 | 4 | 12 | 21 | 0.57 | 0.31 |
| 13 | 93 | 77 | 9 | 7 | 0 | 7 | 16 | 0.44 | 0.21 |
| 14 | 88 | 63 | 6 | 18 | 1 | 18 | 25 | 0.72 | 0.40 |
| 15 | 73 | 47 | 13 | 9 | 4 | 9 | 26 | 0.35 | 0.55 |
| 16 | 67 | 46 | 19 | 2 | 0 | 2 | 21 | 0.10 | 0.46 |
| 17 | 79 | 67 | 9 | 3 | 0 | 3 | 12 | 0.25 | 0.18 |
| 18 | 100 | 76 | 13 | 11 | 0 | 11 | 24 | 0.46 | 0.32 |
| 19 | 93 | 75 | 9 | 8 | 1 | 8 | 18 | 0.44 | 0.24 |
| 20 | 90 | 66 | 17 | 7 | 0 | 7 | 24 | 0.29 | 0.36 |
| 21 | | | | | | | | | |
| 22 | | | | | | | | | |
| 23 | | | | | | | | | |
| 24 | | | | | | | | | |
| 25 | | | | | | | | | |
| 26 | | | | | | | | | |
| 27 | | | | | | | | | |
| 28 | | | | | | | | | |
| 29 | | | | | | | | | |
| 30 | | | | | | | | | |
| 31 | | | | | | | | | |
| 32 | | | | | | | | | |
| 33 | | | | | | | | | |
| 34 | | | | | | | | | |
| 35 | | | | | | | | | |
| 36 | | | | | | | | | |
| 37 | | | | | | | | | |
| 38 | | | | | | | | | |
| 39 | | | | | | | | | |
| 40 | | | | | | | | | |
| TOTAL | 1567 | 1196 | 214 | 134 | 23 | 134 | 371 | 6.52 | 6.64 |
| AVERAGE | 78.4 | 59.8 | 10.7 | 6.7 | 1.2 | 6.7 | 18.6 | 0.33 | 0.33 |
| SEM | 3.5 | 3.2 | 1.0 | 1.1 | 0.3 | 1.1 | 1.4 | 0.04 | 0.03 |

#### p38 Inhibited

| # | TOTAL NUMBER OF CELLS: Embryo | Double Negative | ICM: NANOG +ve | ICM: GATA4 +ve | ICM: NANOG/GATA4 double +ve | ICM: PrE | ICM: Total | Ratio of PrE/Total ICM | Ratio of Inner/Outer cells |
|---|---|---|---|---|---|---|---|---|---|
| 1 | 79 | 68 | 5 | 6 | 0 | 6 | 11 | 0.55 | 0.16 |
| 2 | 57 | 43 | 12 | 2 | 0 | 2 | 14 | 0.14 | 0.33 |
| 3 | 63 | 42 | 15 | 5 | 1 | 5 | 21 | 0.24 | 0.50 |
| 4 | 66 | 57 | 9 | 0 | 0 | 0 | 9 | 0.00 | 0.16 |
| 5 | 60 | 45 | 12 | 3 | 0 | 3 | 15 | 0.20 | 0.33 |
| 6 | 81 | 73 | 2 | 6 | 0 | 6 | 8 | 0.75 | 0.11 |
| 7 | 95 | 79 | 9 | 6 | 1 | 6 | 16 | 0.38 | 0.20 |
| 8 | 59 | 46 | 13 | 0 | 0 | 0 | 13 | 0.00 | 0.28 |
| 9 | 13 | 8 | 5 | 0 | 0 | 0 | 5 | 0.00 | 0.63 |
| 10 | 76 | 68 | 3 | 5 | 0 | 5 | 8 | 0.63 | 0.12 |
| 11 | 28 | 11 | 17 | 0 | 0 | 0 | 17 | 0.00 | 1.55 |
| 12 | 96 | 84 | 8 | 4 | 0 | 4 | 12 | 0.33 | 0.14 |
| 13 | 76 | 61 | 9 | 4 | 2 | 4 | 15 | 0.27 | 0.25 |
| 14 | 71 | 55 | 16 | 0 | 0 | 0 | 16 | 0.00 | 0.29 |
| 15 | 45 | 37 | 8 | 0 | 0 | 0 | 8 | 0.00 | 0.22 |
| 16 | 58 | 44 | 14 | 0 | 0 | 0 | 14 | 0.00 | 0.32 |
| 17 | 85 | 72 | 13 | 0 | 0 | 0 | 13 | 0.00 | 0.18 |
| 18 | 60 | 47 | 11 | 2 | 0 | 2 | 13 | 0.15 | 0.28 |
| 19 | 80 | 71 | 6 | 3 | 0 | 3 | 9 | 0.33 | 0.13 |
| 20 | 63 | 55 | 7 | 1 | 0 | 1 | 8 | 0.13 | 0.15 |
| 21 | 62 | 49 | 5 | 8 | 0 | 8 | 13 | 0.62 | 0.27 |
| 22 | 22 | 19 | 3 | 0 | 0 | 0 | 3 | 0.00 | 0.16 |
| 23 | | | | | | | | | |
| 24 | | | | | | | | | |
| 25 | | | | | | | | | |
| 26 | | | | | | | | | |
| 27 | | | | | | | | | |
| 28 | | | | | | | | | |
| 29 | | | | | | | | | |
| 30 | | | | | | | | | |
| 31 | | | | | | | | | |
| 32 | | | | | | | | | |
| 33 | | | | | | | | | |
| 34 | | | | | | | | | |
| 35 | | | | | | | | | |
| 36 | | | | | | | | | |
| 37 | | | | | | | | | |
| 38 | | | | | | | | | |
| 39 | | | | | | | | | |
| 40 | | | | | | | | | |
| TOTAL | 1395 | 1134 | 202 | 55 | 4 | 55 | 261 | 4.70 | 6.73 |
| AVERAGE | 63.4 | 51.5 | 9.2 | 2.5 | 0.2 | 2.5 | 11.9 | 0.21 | 0.31 |
| SEM | 4.6 | 4.4 | 0.9 | 0.6 | 0.1 | 0.6 | 0.9 | 0.05 | 0.06 |

### KSOM+AA + 10mM NAC

#### DMSO

| # | TOTAL NUMBER OF CELLS: Embryo | Double Negative | ICM: NANOG +ve | ICM: GATA4 +ve | ICM: NANOG/GATA4 double +ve | ICM: PrE | ICM: Total | Ratio of PrE/Total ICM | Ratio of Inner/Outer cells |
|---|---|---|---|---|---|---|---|---|---|
| 1 | 84 | 61 | 17 | 6 | 0 | 6 | 23 | 0.26 | 0.38 |
| 2 | 64 | 54 | 9 | 1 | 0 | 1 | 10 | 0.10 | 0.19 |
| 3 | 66 | 41 | 21 | 3 | 1 | 3 | 25 | 0.12 | 0.61 |
| 4 | 66 | 40 | 21 | 1 | 4 | 1 | 26 | 0.04 | 0.65 |
| 5 | 95 | 67 | 16 | 12 | 0 | 12 | 28 | 0.43 | 0.42 |
| 6 | 58 | 38 | 16 | 3 | 1 | 3 | 20 | 0.15 | 0.53 |
| 7 | 82 | 65 | 7 | 10 | 0 | 10 | 17 | 0.59 | 0.26 |
| 8 | 92 | 76 | 6 | 10 | 0 | 10 | 16 | 0.63 | 0.21 |
| 9 | 45 | 35 | 5 | 4 | 1 | 4 | 10 | 0.40 | 0.29 |
| 10 | 70 | 55 | 7 | 7 | 1 | 7 | 15 | 0.47 | 0.27 |
| 11 | 68 | 57 | 7 | 4 | 0 | 4 | 11 | 0.36 | 0.19 |
| 12 | 57 | 45 | 7 | 3 | 2 | 3 | 12 | 0.25 | 0.27 |
| 13 | 91 | 75 | 9 | 7 | 0 | 7 | 16 | 0.44 | 0.21 |
| 14 | 93 | 78 | 8 | 6 | 1 | 6 | 15 | 0.40 | 0.19 |
| 15 | 84 | 64 | 8 | 11 | 1 | 11 | 20 | 0.55 | 0.31 |
| 16 | 85 | 65 | 10 | 8 | 2 | 8 | 20 | 0.40 | 0.31 |
| 17 | 69 | 59 | 10 | 0 | 0 | 0 | 10 | 0.00 | 0.17 |
| 18 | 71 | 56 | 7 | 6 | 2 | 6 | 15 | 0.40 | 0.27 |
| 19 | 94 | 72 | 11 | 11 | 0 | 11 | 22 | 0.50 | 0.31 |
| 20 | | | | | | | | | |
| TOTAL | 1434 | 1103 | 202 | 113 | 16 | 113 | 331 | 6.48 | 6.03 |
| AVERAGE | 75.5 | 58.1 | 10.6 | 5.9 | 0.8 | 5.9 | 17.4 | 0.34 | 0.32 |
| SEM | 3.4 | 3.0 | 1.2 | 0.8 | 0.2 | 0.8 | 1.3 | 0.04 | 0.03 |

#### p38 Inhibited

| # | TOTAL NUMBER OF CELLS: Embryo | Double Negative | ICM: NANOG +ve | ICM: GATA4 +ve | ICM: NANOG/GATA4 double +ve | ICM: PrE | ICM: Total | Ratio of PrE/Total ICM | Ratio of Inner/Outer cells |
|---|---|---|---|---|---|---|---|---|---|
| 1 | 52 | 30 | 22 | 0 | 0 | 0 | 22 | 0.00 | 0.73 |
| 2 | 51 | 32 | 15 | 4 | 0 | 4 | 19 | 0.21 | 0.59 |
| 3 | 67 | 48 | 18 | 1 | 0 | 1 | 19 | 0.05 | 0.40 |
| 4 | 58 | 52 | 6 | 0 | 0 | 0 | 6 | 0.00 | 0.12 |
| 5 | 55 | 41 | 11 | 1 | 2 | 1 | 14 | 0.07 | 0.34 |
| 6 | 77 | 66 | 4 | 7 | 0 | 7 | 11 | 0.64 | 0.17 |
| 7 | 84 | 71 | 7 | 2 | 4 | 2 | 13 | 0.15 | 0.18 |
| 8 | 61 | 45 | 13 | 3 | 0 | 3 | 16 | 0.19 | 0.36 |
| 9 | 65 | 48 | 12 | 5 | 0 | 5 | 17 | 0.29 | 0.35 |
| 10 | 25 | 19 | 6 | 0 | 0 | 0 | 6 | 0.00 | 0.32 |
| 11 | 53 | 44 | 5 | 3 | 1 | 3 | 9 | 0.33 | 0.20 |
| 12 | 63 | 46 | 12 | 3 | 2 | 3 | 17 | 0.18 | 0.37 |
| 13 | 97 | 80 | 9 | 8 | 0 | 8 | 17 | 0.47 | 0.21 |
| 14 | 60 | 49 | 11 | 0 | 0 | 0 | 11 | 0.00 | 0.22 |
| 15 | 81 | 66 | 11 | 4 | 0 | 4 | 15 | 0.27 | 0.23 |
| 16 | 74 | 57 | 13 | 4 | 0 | 4 | 17 | 0.24 | 0.30 |
| 17 | 67 | 58 | 9 | 0 | 0 | 0 | 9 | 0.00 | 0.16 |
| 18 | 81 | 58 | 11 | 12 | 0 | 12 | 23 | 0.52 | 0.40 |
| 19 | 74 | 65 | 7 | 1 | 1 | 1 | 9 | 0.11 | 0.14 |
| 20 | 106 | 90 | 9 | 7 | 0 | 7 | 16 | 0.44 | 0.18 |
| TOTAL | 1351 | 1065 | 211 | 65 | 10 | 65 | 286 | 4.16 | 5.96 |
| AVERAGE | 67.6 | 53.3 | 10.6 | 3.3 | 0.5 | 3.3 | 14.3 | 0.21 | 0.30 |
| SEM | 4.0 | 3.8 | 1.0 | 0.7 | 0.2 | 0.7 | 1.1 | 0.04 | 0.03 |

Supplementary table S3

### KSOM

#### DMSO

| # | TOTAL NUMBER OF CELLS | | | | | | Ratio of NANOG-GATA6 double +ve/Total ICM |
|---|---|---|---|---|---|---|---|
| | | | ICM | | | | |
| | Embryo | Double Negative | NANOG +ve | NANOG-GATA6 double +ve | PrE | Total | |
| 1 | 75 | 63 | 4 | 6 | 5 | 12 | 0.50 |
| 2 | 65 | 51 | 12 | 9 | 2 | 14 | 0.64 |
| 3 | 70 | 60 | 5 | 4 | 3 | 10 | 0.40 |
| 4 | 65 | 52 | 6 | 11 | 2 | 13 | 0.85 |
| 5 | 70 | 52 | 8 | 4 | 10 | 18 | 0.22 |
| 6 | 97 | 78 | 11 | 8 | 5 | 19 | 0.42 |
| 7 | 89 | 73 | 9 | 4 | 7 | 16 | 0.25 |
| 8 | 82 | 67 | 11 | 6 | 2 | 15 | 0.40 |
| 9 | 92 | 80 | 10 | 3 | 1 | 12 | 0.25 |
| 10 | 82 | 55 | 15 | 8 | 10 | 27 | 0.30 |
| 11 | 87 | 60 | 24 | 23 | 3 | 27 | 0.85 |
| 12 | 88 | 65 | 9 | 7 | 11 | 23 | 0.30 |
| 13 | | | | | | | |
| 14 | | | | | | | |
| 15 | | | | | | | |
| **TOTAL** | **962** | **756** | **124** | **93** | **61** | **206** | 5.38 |
| **AVERAGE** | **80.2** | **63.0** | **10.3** | **7.8** | **5.1** | **17.2** | 0.45 |
| **SEM** | **3.1** | **2.9** | **1.5** | **1.5** | **1.0** | **1.7** | 0.06 |

#### p38 Inhibited

| # | TOTAL NUMBER OF CELLS | | | | | | Ratio of NANOG-GATA6 double +ve/Total ICM |
|---|---|---|---|---|---|---|---|
| | | | ICM | | | | |
| | Embryo | Double Negative | NANOG +ve | NANOG-GATA6 double +ve | PrE | Total | |
| 1 | 70 | 65 | 3 | 5 | 0 | 5 | 1.00 |
| 2 | 37 | 35 | 2 | 2 | 0 | 2 | 1.00 |
| 3 | 75 | 67 | 6 | 5 | 2 | 8 | 0.63 |
| 4 | 38 | 21 | 17 | 17 | 0 | 17 | 1.00 |
| 5 | 38 | 29 | 9 | 9 | 0 | 9 | 1.00 |
| 6 | 90 | 84 | 5 | 2 | 1 | 6 | 0.33 |
| 7 | 45 | 37 | 8 | 8 | 0 | 8 | 1.00 |
| 8 | 27 | 16 | 11 | 11 | 0 | 11 | 1.00 |
| 9 | 40 | 35 | 5 | 5 | 0 | 5 | 1.00 |
| 10 | 37 | 30 | 5 | 7 | 0 | 7 | 1.00 |
| 11 | 52 | 43 | 9 | 9 | 0 | 9 | 1.00 |
| 12 | 45 | 34 | 11 | 11 | 0 | 11 | 1.00 |
| 13 | | | | | | | |
| 14 | | | | | | | |
| 15 | | | | | | | |
| **TOTAL** | **594** | **496** | **91** | **91** | **3** | **98** | 10.96 |
| **AVERAGE** | **49.5** | **41.3** | **7.6** | **7.6** | **0.3** | **8.2** | 0.91 |
| **SEM** | **5.5** | **5.9** | **1.2** | **1.2** | **0.2** | **1.1** | 0.06 |

### KSOM with amino acids

#### DMSO

| # | TOTAL NUMBER OF CELLS | | | | | | Ratio of NANOG-GATA6 double +ve/Total ICM |
|---|---|---|---|---|---|---|---|
| | | | ICM | | | | |
| | Embryo | Double Negative | NANOG +ve | NANOG-GATA6 double +ve | PrE | Total | |
| 1 | 84 | 60 | 9 | 0 | 15 | 24 | 0.00 |
| 2 | 86 | 62 | 14 | 11 | 9 | 24 | 0.46 |
| 3 | 89 | 70 | 13 | 4 | 6 | 19 | 0.21 |
| 4 | 72 | 58 | 11 | 8 | 1 | 14 | 0.57 |
| 5 | 58 | 48 | 6 | 2 | 4 | 10 | 0.20 |
| 6 | 73 | 52 | 6 | 2 | 15 | 21 | 0.10 |
| 7 | 85 | 63 | 15 | 7 | 5 | 22 | 0.32 |
| 8 | 90 | 67 | 7 | 2 | 15 | 23 | 0.09 |
| 9 | 85 | 67 | 9 | 6 | 7 | 18 | 0.33 |
| 10 | 117 | 98 | 11 | 4 | 7 | 19 | 0.21 |
| 11 | 84 | 74 | 6 | 2 | 4 | 10 | 0.20 |
| 12 | 88 | 69 | 10 | 3 | 8 | 19 | 0.16 |
| 13 | 107 | 89 | 11 | 2 | 7 | 18 | 0.11 |
| 14 | 98 | 68 | 12 | 3 | 16 | 30 | 0.10 |
| 15 | 94 | 81 | 11 | 9 | 1 | 13 | 0.69 |
| 16 | 90 | 67 | 16 | 6 | 7 | 23 | 0.26 |
| 17 | 80 | 60 | 10 | 7 | 8 | 20 | 0.35 |
| 18 | 79 | 57 | 8 | 3 | 12 | 22 | 0.14 |
| 19 | 93 | 71 | 19 | 14 | 3 | 22 | 0.64 |
| 20 | 66 | 45 | 18 | 4 | 3 | 21 | 0.19 |
| 21 | 101 | 74 | 14 | 2 | 11 | 27 | 0.07 |
| 22 | 72 | 55 | 12 | 3 | 5 | 17 | 0.18 |
| 23 | 83 | 57 | 15 | 7 | 9 | 26 | 0.27 |
| 24 | 71 | 40 | 21 | 14 | 5 | 31 | 0.45 |
| 25 | 92 | 73 | 8 | 5 | 7 | 19 | 0.26 |
| 26 | 78 | 58 | 11 | 8 | 3 | 20 | 0.40 |
| 27 | 80 | 60 | 13 | 0 | 7 | 20 | 0.00 |
| 28 | 77 | 56 | 10 | 2 | 11 | 21 | 0.10 |
| 29 | 60 | 40 | 20 | 11 | 0 | 20 | 0.55 |
| 30 | 78 | 69 | 6 | 0 | 3 | 9 | 0.00 |
| 31 | 95 | 75 | 8 | 6 | 9 | 20 | 0.30 |
| 32 | 106 | 88 | 13 | 6 | 4 | 18 | 0.33 |
| 33 | 80 | 62 | 8 | 2 | 8 | 18 | 0.11 |
| 34 | 111 | 82 | 12 | 0 | 17 | 29 | 0.00 |
| 35 | 110 | 88 | 10 | 1 | 12 | 22 | 0.05 |
| 36 | 76 | 59 | 10 | 6 | 7 | 17 | 0.35 |
| 37 | 96 | 76 | 10 | 3 | 10 | 20 | 0.15 |
| 38 | | | | | | | |
| 39 | | | | | | | |

#### p38 Inhibited

| # | TOTAL NUMBER OF CELLS | | | | | | Ratio of NANOG-GATA6 double +ve/Total ICM |
|---|---|---|---|---|---|---|---|
| | | | ICM | | | | |
| | Embryo | Double Negative | NANOG +ve | NANOG-GATA6 double +ve | PrE | Total | |
| 1 | 75 | 65 | 7 | 1 | 3 | 10 | 0.10 |
| 2 | 62 | 43 | 16 | 4 | 3 | 19 | 0.21 |
| 3 | 61 | 47 | 12 | 7 | 2 | 14 | 0.50 |
| 4 | 79 | 66 | 9 | 2 | 4 | 13 | 0.15 |
| 5 | 23 | 18 | 5 | 5 | 0 | 5 | 1.00 |
| 6 | 68 | 54 | 9 | 2 | 4 | 14 | 0.14 |
| 7 | 68 | 59 | 5 | 1 | 4 | 9 | 0.11 |
| 8 | 57 | 43 | 9 | 2 | 5 | 14 | 0.14 |
| 9 | 63 | 50 | 13 | 4 | 0 | 13 | 0.31 |
| 10 | 64 | 59 | 5 | 5 | 0 | 5 | 1.00 |
| 11 | 66 | 55 | 10 | 9 | 1 | 11 | 0.82 |
| 12 | 52 | 44 | 4 | 2 | 3 | 8 | 0.25 |
| 13 | 68 | 61 | 7 | 6 | 0 | 7 | 0.86 |
| 14 | 66 | 50 | 15 | 11 | 1 | 16 | 0.69 |
| 15 | 60 | 49 | 7 | 8 | 2 | 11 | 0.73 |
| 16 | 49 | 37 | 12 | 3 | 0 | 12 | 0.25 |
| 17 | 57 | 50 | 7 | 1 | 0 | 7 | 0.14 |
| 18 | 85 | 60 | 16 | 4 | 8 | 25 | 0.16 |
| 19 | 63 | 54 | 8 | 8 | 0 | 9 | 0.89 |
| 20 | 71 | 60 | 11 | 7 | 0 | 11 | 0.64 |
| 21 | 53 | 34 | 18 | 10 | 0 | 19 | 0.53 |
| 22 | 51 | 48 | 3 | 1 | 0 | 3 | 0.33 |
| 23 | 49 | 39 | 9 | 6 | 1 | 10 | 0.60 |
| 24 | 35 | 27 | 7 | 7 | 1 | 8 | 0.88 |
| 25 | 45 | 28 | 17 | 9 | 0 | 17 | 0.53 |
| 26 | 71 | 59 | 8 | 4 | 1 | 12 | 0.33 |
| 27 | 55 | 49 | 5 | 2 | 1 | 6 | 0.33 |
| 28 | 59 | 29 | 30 | 27 | 0 | 30 | 0.90 |
| 29 | 43 | 24 | 19 | 17 | 0 | 19 | 0.89 |
| 30 | 52 | 44 | 5 | 2 | 2 | 8 | 0.25 |
| 31 | 55 | 34 | 20 | 8 | 0 | 21 | 0.38 |
| 32 | 66 | 56 | 10 | 2 | 0 | 10 | 0.20 |
| 33 | 65 | 50 | 11 | 9 | 4 | 15 | 0.60 |
| 34 | 62 | 49 | 11 | 6 | 2 | 13 | 0.46 |
| 35 | 48 | 36 | 12 | 9 | 0 | 12 | 0.75 |
| 36 | 85 | 69 | 6 | 0 | 10 | 16 | 0.00 |
| 37 | 73 | 56 | 14 | 9 | 1 | 17 | 0.53 |
| 38 | 76 | 54 | 11 | 2 | 11 | 22 | 0.09 |
| 39 | | | | | | | |

| | | | | | | | |
|---|---|---|---|---|---|---|---|
| 40 | | | | | | | |
| TOTAL | **3184** | **2438** | **423** | **175** | **281** | **746** | 8.89 |
| AVERAGE | **86.1** | **65.9** | **11.4** | **4.7** | **7.6** | **20.2** | 0.24 |
| SEM | **2.3** | **2.2** | **0.6** | **0.6** | **0.7** | **0.8** | 0.03 |

| | | | | | | | |
|---|---|---|---|---|---|---|---|
| 40 | | | | | | | |
| TOTAL | **2300** | **1809** | **403** | **222** | **74** | **491** | 17.68 |
| AVERAGE | **60.5** | **47.6** | **10.6** | **5.8** | **1.9** | **12.9** | 0.47 |
| SEM | **2.1** | **2.0** | **0.9** | **0.8** | **0.4** | **0.9** | 0.05 |

Supplementary table S3 (contd.)

### KSOM + 1mM NAC

#### DMSO

| # | TOTAL NUMBER OF CELLS | | | | | | Ratio of NANOG-GATA6 double +ve/Total ICM |
|---|---|---|---|---|---|---|---|
| | Embryo | Double Negative | ICM | | | | |
| | | | NANOG +ve | NANOG-GATA6 double +ve | PrE | Total | |
| 1 | 97 | 75 | 13 | 4 | 8 | 22 | 0.18 |
| 2 | 73 | 51 | 17 | 17 | 2 | 22 | 0.77 |
| 3 | 70 | 43 | 14 | 5 | 10 | 27 | 0.19 |
| 4 | 91 | 68 | 13 | 4 | 10 | 23 | 0.17 |
| 5 | 62 | 50 | 7 | 5 | 2 | 12 | 0.42 |
| 6 | 75 | 47 | 18 | 7 | 9 | 28 | 0.25 |
| 7 | 91 | 73 | 10 | 10 | 7 | 18 | 0.56 |
| 8 | 64 | 52 | 6 | 6 | 6 | 12 | 0.50 |
| 9 | 84 | 71 | 8 | 3 | 4 | 13 | 0.23 |
| 10 | 77 | 58 | 11 | 3 | 8 | 19 | 0.16 |
| 11 | 84 | 70 | 9 | 2 | 5 | 14 | 0.14 |
| 12 | | | | | | | |
| **TOTAL** | **868** | **658** | **126** | **66** | **71** | **210** | 3.57 |
| **AVERAGE** | **78.9** | **59.8** | **11.5** | **6.0** | **6.5** | **19.1** | 0.32 |
| **SEM** | **3.5** | **3.5** | **1.2** | **1.3** | **0.9** | **1.8** | 0.06 |

#### p38 Inhibited

| # | TOTAL NUMBER OF CELLS | | | | | | Ratio of NANOG-GATA6 double +ve/Total ICM |
|---|---|---|---|---|---|---|---|
| | Embryo | Double Negative | ICM | | | | |
| | | | NANOG +ve | NANOG-GATA6 double +ve | PrE | Total | |
| 1 | 78 | 67 | 11 | 8 | 0 | 11 | 0.73 |
| 2 | 83 | 81 | 2 | 0 | 0 | 2 | 0.00 |
| 3 | 81 | 70 | 7 | 2 | 4 | 11 | 0.18 |
| 4 | 47 | 46 | 1 | 0 | 0 | 1 | 0.00 |
| 5 | 45 | 38 | 7 | 7 | 0 | 7 | 1.00 |
| 6 | 75 | 67 | 4 | 0 | 4 | 8 | 0.00 |
| 7 | 65 | 61 | 2 | 0 | 2 | 4 | 0.00 |
| 8 | 60 | 51 | 9 | 6 | 0 | 9 | 0.67 |
| 9 | 70 | 63 | 4 | 2 | 3 | 7 | 0.29 |
| 10 | 60 | 51 | 9 | 7 | 0 | 9 | 0.78 |
| 11 | | | | | | | |
| 12 | | | | | | | |
| **TOTAL** | **664** | **595** | **56** | **32** | **13** | **69** | 3.64 |
| **AVERAGE** | **66.4** | **59.5** | **5.6** | **3.2** | **1.3** | **6.9** | 0.36 |
| **SEM** | **4.3** | **4.1** | **1.1** | **1.1** | **0.6** | **1.1** | 0.12 |

### KSOM + 10mM NAC

#### DMSO

| # | TOTAL NUMBER OF CELLS | | | | | | Ratio of NANOG-GATA6 double +ve/Total ICM |
|---|---|---|---|---|---|---|---|
| | Embryo | Double Negative | ICM | | | | |
| | | | NANOG +ve | NANOG-GATA6 double +ve | PrE | Total | |
| 1 | 103 | 83 | 10 | 2 | 8 | 20 | 0.10 |
| 2 | 81 | 60 | 10 | 3 | 11 | 21 | 0.14 |
| 3 | 82 | 61 | 9 | 4 | 11 | 21 | 0.19 |
| 4 | 95 | 78 | 7 | 2 | 10 | 17 | 0.12 |
| 5 | 83 | 72 | 8 | 0 | 3 | 11 | 0.00 |
| 6 | 93 | 75 | 9 | 3 | 9 | 18 | 0.17 |
| 7 | 83 | 66 | 9 | 1 | 8 | 17 | 0.06 |
| 8 | 102 | 86 | 7 | 0 | 9 | 16 | 0.00 |
| 9 | 72 | 48 | 12 | 1 | 12 | 24 | 0.04 |
| 10 | | | | | | | |
| **TOTAL** | **794** | **629** | **81** | **16** | **81** | **165** | 0.82 |
| **AVERAGE** | **88.2** | **69.9** | **9.0** | **1.8** | **9.0** | **18.3** | 0.09 |
| **SEM** | **3.5** | **4.1** | **0.5** | **0.5** | **0.9** | **1.2** | 0.02 |

#### p38 Inhibited

| # | TOTAL NUMBER OF CELLS | | | | | | Ratio of NANOG-GATA6 double +ve/Total ICM |
|---|---|---|---|---|---|---|---|
| | Embryo | Double Negative | ICM | | | | |
| | | | NANOG +ve | NANOG-GATA6 double +ve | PrE | Total | |
| 1 | 68 | 55 | 9 | 3 | 4 | 13 | 0.23 |
| 2 | 90 | 74 | 8 | 1 | 8 | 16 | 0.06 |
| 3 | 69 | 61 | 8 | 6 | 0 | 8 | 0.75 |
| 4 | 78 | 59 | 13 | 7 | 6 | 19 | 0.37 |
| 5 | 62 | 52 | 10 | 7 | 0 | 10 | 0.70 |
| 6 | 47 | 39 | 6 | 4 | 2 | 8 | 0.50 |
| 7 | 81 | 67 | 8 | 2 | 6 | 14 | 0.14 |
| 8 | 72 | 64 | 7 | 7 | 1 | 8 | 0.88 |
| 9 | 52 | 45 | 5 | 3 | 2 | 7 | 0.43 |
| 10 | 36 | 25 | 8 | 3 | 3 | 11 | 0.27 |
| **TOTAL** | **655** | **541** | **82** | **43** | **32** | **114** | 4.33 |
| **AVERAGE** | **65.5** | **54.1** | **8.2** | **4.3** | **3.2** | **11.4** | 0.43 |
| **SEM** | **5.2** | **4.6** | **0.7** | **0.7** | **0.9** | **1.3** | 0.09 |

Supplementary table S3 (contd. 2)

### KSOM+AA + 1mM NAC

#### DMSO

| # | TOTAL NUMBER OF CELLS | | | | | | Ratio of NANOG-GATA6 double +ve/Total ICM |
|---|---|---|---|---|---|---|---|
| | | | ICM | | | | |
| | Embryo | Double Negative | NANOG +ve | NANOG-GATA6 double +ve | PrE | Total | |
| 1 | 97 | 75 | 12 | 3 | 9 | 22 | 0.14 |
| 2 | 87 | 77 | 7 | 9 | 1 | 10 | 0.90 |
| 3 | 49 | 33 | 10 | 13 | 3 | 16 | 0.81 |
| 4 | 45 | 30 | 11 | 13 | 1 | 15 | 0.87 |
| 5 | 68 | 56 | 9 | 9 | 2 | 12 | 0.75 |
| 6 | 78 | 50 | 17 | 6 | 11 | 28 | 0.21 |
| 7 | 78 | 62 | 10 | 3 | 5 | 16 | 0.19 |
| 8 | 56 | 53 | 3 | 1 | 0 | 3 | 0.33 |
| 9 | 68 | 45 | 16 | 12 | 5 | 23 | 0.52 |
| 10 | 91 | 73 | 6 | 0 | 12 | 18 | 0.00 |
| 11 | 78 | 57 | 13 | 6 | 8 | 21 | 0.29 |
| 12 | 89 | 68 | 5 | 4 | 12 | 21 | 0.19 |
| 13 | 93 | 77 | 9 | 2 | 7 | 16 | 0.13 |
| 14 | 88 | 63 | 6 | 3 | 18 | 25 | 0.12 |
| 15 | 73 | 47 | 13 | 11 | 9 | 26 | 0.42 |
| 16 | 67 | 46 | 19 | 12 | 2 | 21 | 0.57 |
| 17 | 79 | 67 | 9 | 3 | 3 | 12 | 0.25 |
| 18 | 100 | 76 | 13 | 4 | 11 | 24 | 0.17 |
| 19 | 93 | 75 | 9 | 6 | 8 | 18 | 0.33 |
| 20 | 90 | 66 | 17 | 10 | 7 | 24 | 0.42 |
| 21 | | | | | | | |
| 22 | | | | | | | |
| 23 | | | | | | | |
| 24 | | | | | | | |
| 25 | | | | | | | |
| **TOTAL** | **1567** | **1196** | **214** | **130** | **134** | **371** | 7.60 |
| **AVERAGE** | **78.4** | **59.8** | **10.7** | **6.5** | **6.7** | **18.6** | 0.38 |
| **SEM** | **3.5** | **3.2** | **1.0** | **1.0** | **1.1** | **1.4** | 0.06 |

#### p38 Inhibited

| # | TOTAL NUMBER OF CELLS | | | | | | Ratio of NANOG-GATA6 double +ve/Total ICM |
|---|---|---|---|---|---|---|---|
| | | | ICM | | | | |
| | Embryo | Double Negative | NANOG +ve | NANOG-GATA6 double +ve | PrE | Total | |
| 1 | 79 | 68 | 5 | 0 | 6 | 11 | 0.00 |
| 2 | 57 | 43 | 12 | 2 | 2 | 14 | 0.14 |
| 3 | 63 | 42 | 15 | 3 | 5 | 21 | 0.14 |
| 4 | 66 | 57 | 9 | 4 | 0 | 9 | 0.44 |
| 5 | 60 | 45 | 12 | 5 | 3 | 15 | 0.33 |
| 6 | 81 | 73 | 2 | 0 | 6 | 8 | 0.00 |
| 7 | 95 | 79 | 9 | 1 | 6 | 16 | 0.06 |
| 8 | 59 | 46 | 13 | 5 | 0 | 13 | 0.38 |
| 9 | 13 | 8 | 5 | 0 | 0 | 5 | 0.00 |
| 10 | 76 | 68 | 3 | 0 | 5 | 8 | 0.00 |
| 11 | 28 | 11 | 17 | 14 | 0 | 17 | 0.82 |
| 12 | 96 | 84 | 8 | 3 | 4 | 12 | 0.25 |
| 13 | 76 | 61 | 9 | 2 | 4 | 15 | 0.13 |
| 14 | 71 | 55 | 16 | 8 | 0 | 16 | 0.50 |
| 15 | 45 | 37 | 8 | 6 | 0 | 8 | 0.75 |
| 16 | 58 | 44 | 14 | 12 | 0 | 14 | 0.86 |
| 17 | 85 | 72 | 13 | 9 | 0 | 13 | 0.69 |
| 18 | 60 | 47 | 11 | 9 | 2 | 13 | 0.69 |
| 19 | 80 | 71 | 6 | 3 | 3 | 9 | 0.33 |
| 20 | 63 | 55 | 7 | 5 | 1 | 8 | 0.63 |
| 21 | 62 | 49 | 5 | 1 | 8 | 13 | 0.08 |
| 22 | 22 | 19 | 3 | 2 | 0 | 3 | 0.67 |
| 23 | | | | | | | |
| 24 | | | | | | | |
| 25 | | | | | | | |
| **TOTAL** | **1395** | **1134** | **202** | **94** | **55** | **261** | 7.91 |
| **AVERAGE** | **63.4** | **51.5** | **9.2** | **4.3** | **2.5** | **11.9** | 0.36 |
| **SEM** | **4.6** | **4.4** | **0.9** | **0.8** | **0.6** | **0.9** | 0.06 |

### KSOM+AA + 10mM NAC

#### DMSO

| # | TOTAL NUMBER OF CELLS | | | | | | Ratio of NANOG-GATA6 double +ve/Total ICM |
|---|---|---|---|---|---|---|---|
| | | | ICM | | | | |
| | Embryo | Double Negative | NANOG +ve | NANOG-GATA6 double +ve | PrE | Total | |
| 1 | 84 | 61 | 17 | 4 | 6 | 23 | 0.17 |
| 2 | 64 | 54 | 9 | 4 | 1 | 10 | 0.40 |
| 3 | 66 | 41 | 21 | 9 | 3 | 25 | 0.36 |
| 4 | 66 | 40 | 21 | 8 | 1 | 26 | 0.31 |
| 5 | 95 | 67 | 16 | 1 | 12 | 28 | 0.04 |
| 6 | 58 | 38 | 16 | 11 | 3 | 20 | 0.55 |
| 7 | 92 | 76 | 6 | 0 | 10 | 16 | 0.00 |
| 8 | 45 | 35 | 5 | 1 | 4 | 10 | 0.10 |
| 9 | 70 | 55 | 7 | 1 | 7 | 15 | 0.07 |
| 10 | 91 | 75 | 9 | 0 | 7 | 16 | 0.00 |
| 11 | 93 | 78 | 8 | 4 | 6 | 15 | 0.27 |
| 12 | 84 | 64 | 8 | 2 | 11 | 20 | 0.10 |
| 13 | 85 | 65 | 10 | 4 | 8 | 20 | 0.20 |
| 14 | 69 | 59 | 10 | 10 | 0 | 10 | 1.00 |
| 15 | 71 | 56 | 7 | 3 | 6 | 15 | 0.20 |
| 16 | 94 | 72 | 11 | 1 | 11 | 22 | 0.05 |
| 17 | | | | | | | |
| 18 | | | | | | | |
| 19 | | | | | | | |
| 20 | | | | | | | |
| **TOTAL** | **1227** | **936** | **181** | **63** | **96** | **291** | 3.81 |
| **AVERAGE** | **76.7** | **58.5** | **11.3** | **3.9** | **6.0** | **18.2** | 0.24 |
| **SEM** | **3.8** | **3.5** | **1.3** | **0.9** | **0.9** | **1.4** | 0.06 |

#### p38 Inhibited

| # | TOTAL NUMBER OF CELLS | | | | | | Ratio of NANOG-GATA6 double +ve/Total ICM |
|---|---|---|---|---|---|---|---|
| | | | ICM | | | | |
| | Embryo | Double Negative | NANOG +ve | NANOG-GATA6 double +ve | PrE | Total | |
| 1 | 25 | 19 | 6 | 2 | 0 | 6 | 0.33 |
| 2 | 53 | 44 | 5 | 1 | 3 | 9 | 0.11 |
| 3 | 63 | 46 | 12 | 3 | 3 | 17 | 0.18 |
| 4 | 97 | 80 | 9 | 0 | 8 | 17 | 0.00 |
| 5 | 60 | 49 | 11 | 7 | 0 | 11 | 0.64 |
| 6 | 81 | 66 | 11 | 3 | 4 | 15 | 0.20 |
| 7 | 74 | 57 | 13 | 2 | 4 | 17 | 0.12 |
| 8 | 67 | 58 | 9 | 5 | 0 | 9 | 0.56 |
| 9 | 81 | 58 | 11 | 0 | 12 | 23 | 0.00 |
| 10 | 74 | 65 | 7 | 4 | 1 | 9 | 0.44 |
| 11 | 106 | 90 | 9 | 0 | 7 | 16 | 0.00 |
| 12 | | | | | | | |
| 13 | | | | | | | |
| 14 | | | | | | | |
| 15 | | | | | | | |
| 16 | | | | | | | |
| 17 | | | | | | | |
| 18 | | | | | | | |
| 19 | | | | | | | |
| 20 | | | | | | | |
| **TOTAL** | **781** | **632** | **103** | **27** | **42** | **149** | 2.57 |
| **AVERAGE** | **71.0** | **57.5** | **9.4** | **2.5** | **3.8** | **13.5** | 0.23 |
| **SEM** | **6.6** | **5.7** | **0.8** | **0.7** | **1.2** | **1.5** | 0.07 |

**Supplementary table S4 (antibodies used)**

| Sl. No. | Antibody | Type | Target | Host / isotype | Manufacturer | Catalogue number | Dilution used |
|---|---|---|---|---|---|---|---|
| **Immunofluorescence imaging** | | | | | | | |
| 1 | Nanog (eBioMLC-51) | Primary; monoclonal | Mouse | Rat / IgG2a | Thermo Fisher Scientific Inc. (eBioscience™) | 14-5761-80 | 1:200 in PBST (3% BSA) |
| 2 | GATA-4 (H-112) | Primary; polyclonal | Mouse, rat and human | Rabbit / IgG | Santa Cruz Biotechnology, Inc. | sc-9053 | 1:200 in PBST (3% BSA) |
| 3 | GATA-6 | Primary; polyclonal | Human and mouse | Goat / IgG | R&D Systems™ | AF1700 | 1:200 in PBST (3% BSA) |
| 4 | Donkey anti-Rat IgG (H+L) Highly Cross-Adsorbed Secondary Antibody, Alexa Fluor 488 | Secondary; polyclonal | Rat | Donkey / IgG | Thermo Fisher Scientific Inc. | A-21208 | 1:500 in PBST (3% BSA) |
| 5 | Donkey anti-Rabbit IgG (H+L) Highly Cross-Adsorbed Secondary Antibody, Alexa Fluor 555 | Secondary; polyclonal | Rabbit | Donkey / IgG | Thermo Fisher Scientific Inc. | A-31572 | 1:500 in PBST (3% BSA) |
| 6 | Donkey Anti-Goat IgG H&L (Alexa Fluor® 647) | Secondary; polyclonal | Goat | Donkey / IgG | Abcam plc. | ab150131 | 1:500 in PBST (3% BSA) |
| 7 | Donkey Anti-Rabbit IgG H&L (Alexa Fluor® 647) | Secondary; polyclonal | Rabbit | Donkey / IgG | Abcam plc. | ab150075 | 1:500 in PBST (3% BSA) |
| 8 | Goat Anti-Rat IgG H&L (Cy3 ®) preadsorbed | Secondary; polyclonal | Rat | Goat / IgG | Abcam plc. | ab98416 | 1:500 in PBST (3% BSA) |
| **Immunoblotting** | | | | | | | |
| 9 | Phospho-p38 MAPK (Thr180/Tyr182) (28B10) | Primary; monoclonal | Mouse, rat and human | Mouse / IgG1 | Cell Signaling Technology, Inc. | 9216S | 1:500 in 1% milk |
| 10 | GAPDH | Primary; polyclonal | Mouse, rat and human | Rabbit | Merck KGaA (Sigma-Aldrich, Inc.) | G9545 | 1:20,000 in 1% milk |
| 11 | Peroxidase AffiniPure Goat Anti-Mouse IgG (H+L) | Secondary; polyclonal | Mouse | Goat / IgG | Jackson ImmunoResearch Europe Ltd. | 115-035-003 | 1:10,000 in 1% milk |
| 12 | Peroxidase AffiniPure Donkey Anti-Rabbit IgG (H+L) | Secondary; polyclonal | Rabbit | Donkey / IgG | Jackson ImmunoResearch Europe Ltd. | 711-035-152 | 1:10,000 in 1% milk |

**Supplementary table S5 (ANOVA results for figures 4 and 5e & e')**

**Welch's ANOVA test**

| | KSOM | | KSOM+AA | |
|---|---|---|---|---|
| | DMSO | SB220025 (p38i) | DMSO | SB220025 (p38i) |
| **Total cell number** | 0.5568 (ns) | 0.0005 (***) | 0.2422 (ns) | 0.1291 (ns) |
| **Outer cell number** | 0.6594 (ns) | 0.0019 (**) | 0.2643 (ns) | 0.2595 (ns) |
| **Inner cell number** | 0.6266 (ns) | 0.0229 (*) | 0.7817 (ns) | 0.1476 (ns) |
| **Epiblast cell number** | 0.0959 (ns) | 0.1745 (ns) | 0.9893 (ns) | 0.5693 (ns) |
| **PrE cell number** | 0.9329 (ns) | 0.0022 (**) | 0.5677 (ns) | 0.2169 (ns) |
| **PrE to ICM ratio** | 0.6893 (ns) | 0.0015 (**) | 0.6342 (ns) | 0.2126 (ns) |
| **Uncommitted to ICM ratio** | <0.0001 (****) | 0.0002 (***) | 0.1308 (ns) | 0.0348 (*) |

**Supplementary table S6 (oligonucleotide primers used for Q-RT PCR; Fig 3h)**

| Sequences of oligonucleotide primer pairs used for Q-RT PCR | | | |
|---|---|---|---|
| **Sl. No.** | **Gene** | **Forward primer (5'-3')** | **Reverse primer (5'-3')** |
| 1 | H2afz | GCGCAGCCATCCTGGAGTA | CCGATCAGCGATTTGTGGA |
| 2 | Cat | TCCTCGTTCAGGATGTGGTT | TGCGTGTACCACTCTCTCAG |
| 3 | Sod1 | CGGTGAACCAGTTGTGTTGT | GGTCTCCAACATGCCTCTCT |
| 4 | Sod2 | CAAGCACAGCCTCCCAGA | ATCTGCGCGTTAATGTGTGG |

Supplementary Fig. S1 (supplementary to Fig 3 b (fixed))

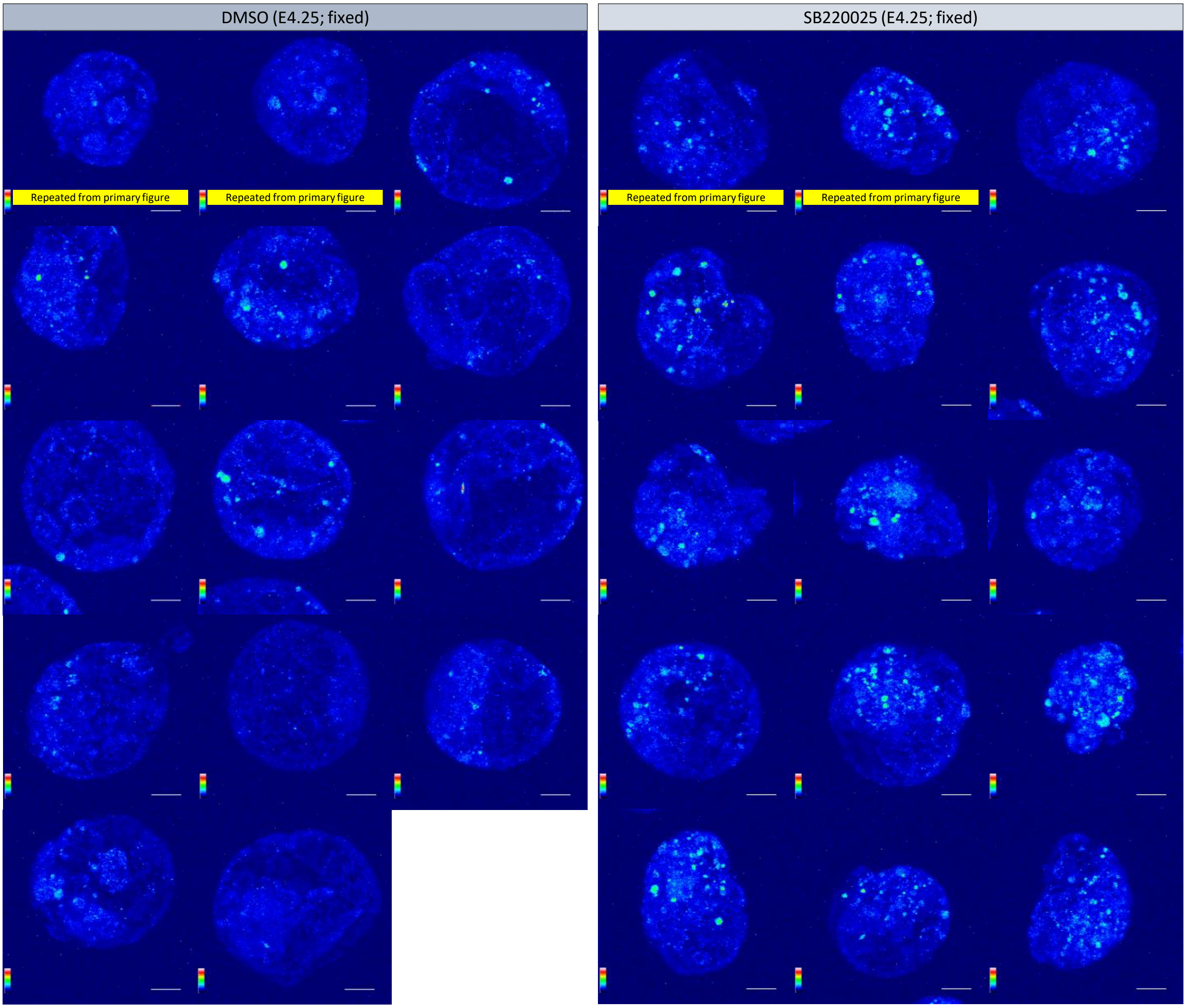


(Supplementary table to Fig 3 c)

Count of ROX-positive foci (maxima) for E4.25

| | KSOM | |
|---|---|---|
| | DMSO | SB220025 |
| | 204 | 442 |
| | 291 | 422 |
| | 314 | 466 |
| | 456 | 462 |
| | 543 | 617 |
| | 423 | 541 |
| | 393 | 589 |
| | 591 | 623 |
| | 385 | 478 |
| | 393 | 520 |
| | 144 | 668 |
| | 461 | 608 |
| | 389 | 640 |
| | 237 | 346 |
| | | 567 |
| Average | 373.14 | 532.60 |
